# Analyses of the core eukaryotic protein subunit of telomerase support extensive adaptation to different evolutionary and life histories in the Metazoa

**DOI:** 10.1101/091124

**Authors:** Alvina G. Lai, Natalia Pouchkina-Stantcheva, Alessia Di Donfrancesco, Gerda Kildisiute, Sounak Sahu, A. Aziz Aboobaker

**Author notes:** Corresponding authors: Alvina G. Lai A. Aziz Aboobaker.

## Abstract

Most animals employ telomerase, which consists of a catalytic subunit known as the telomerase reverse transcriptase (TERT) and an RNA template, to maintain telomere ends. Given the importance of TERT and the apparent importance of telomere biology in core metazoan life history traits like ageing and the control of somatic cell proliferation, we hypothesised that TERT would have patterns of sequence and regulatory evolution reflecting adaptations to diverse evolutionary and life histories across the Animal Kingdom. To test this, we performed a complete investigation of the evolutionary history of TERT across animals. We show that although TERT is almost ubiquitous across Metazoa, it has undergone substantial sequence evolution in canonical motifs. Beyond the known canonical motifs, we also identify and compare regions that are highly variable between lineages, but for which conservation exists within phyla. Recent data have highlighted the importance of alternate splice forms of TERT in non-canonical functions in some animals. Although animals may share some conserved introns, we find that the selection of exons for alternative splicing appears to be highly variable, and regulation by alternative splicing appears to be a very dynamic feature of TERT evolution. We show that even within a closely related group of triclad flatworms, where alternative splicing of TERT was previously correlated with reproductive strategy, we observe highly diverse alternative splicing patterns. Our work establishes that the evolutionary history and structural evolution of TERT involves previously unappreciated levels of change, supporting the view that this core eukaryotic protein has adapted to the requirements of diverse animal life histories.

## Introduction

In linear eukaryotic chromosomes, telomeres function as protective nucleoprotein structures that guard chromosome ends against the action of nucleases, DNA damage signalling and nucleolytic degradation (Muller 1928; Harley et al. 1990; De lange 2005). Telomeric DNA, consisting entirely of guanine-rich tandem repeats, is maintained by the action of the enzyme telomerase. The activity of telomerase was first eluded with the framing of the “end replication problem” by Watson (1972) and Olovnikov (1973), which noted that the lagging strand nature of DNA replication would lead to progressive shortening of linear eukaryotic chromosomes. In humans, cells in culture have a finite replicative lifespan limiting the number of times a cell can divide, and eventually senesce (Hayflick and Moorhead 1961). This has since been shown to be a key mechanism in protecting against cancer and may also contribute to causing a decline in physiological functions and organismal ageing (Harley et al. 1995; Rudolph et al. 1999; Autexier and Lue 2006; Blackburn et al. 2006; Tomás-Loba et al. 2008; de Jesus et al. 2012). While together the shortening of chromosome ends and the observation of the “Hayflick limit” led to the notion that telomeres could act as timers of cellular replicative age, it also raised the question as to how chromosomes maintain their ends across generations, through reproduction, development and during the lifespan of long-lived individuals (Harley et al. 1990; Counter et al. 1992; Vaziri et al. 1994; Bryan et al. 1995). This led to intensive research on both the structure of chromosome ends and a search for the cellular activities that could maintain these structures in light of the “end replication problem”. Eventually, the ribonucleoprotein telomerase was discovered in the ciliated protozoa *Tetrahymena thermophila* (Blackburn and Gall 1978; Szostak and Blackburn 1982; Greider and Blackburn 1985; Greider and Blackburn 1987; Greider and Blackburn 1989). Soon after, the catalytic subunit of this enzyme, telomerase reverse transcriptase (TERT) was also discovered and then studied in human cells (Kilian et al. 1997; Harrington et al. 1997; Meyerson et al. 1997; Nakamura et al. 1997; Lingner et al. 1997; Nugent and Lundblad 1998; Nakayama et al. 1998).

However, despite telomeres and telomerase providing the solution for the existence of linear chromosomes, it is clear that the huge variation in life histories, particularly amongst metazoans, would have different demands on an end chromosome maintenance system. For example, short-lived and long-lived metazoans would have very different needs for telomerase and telomere maintenance. This is exemplified by comparing ourselves with single celled protists, where human life history in an evolutionary context means that signalling from chromosome ends must achieve a balance between allowing growth and preventing inevitable somatic mutations from causing cancer and death before reproductive age (Rubio et al. 2004; Suram and Herbig 2014). Subsequent pleiotropic effects of signalling mechanisms associated with telomeres now appear to be central to the human ageing process and age associated pathologies, including processes that are unconnected to telomere length (Bischoff et al. 2006; Kimura et al. 2008; Steenstrup et al. 2013; Stone et al. 2016). On this basis we would expect to observe telomere biology adaptations to reflect life history specific requirements and therefore variation between different groups. For instance, telomerase levels in the long-lived rodent, the naked-mole rat show stable expression across different life stages (Kim et al. 2011). In long-lived birds, telomerase activity is correlated with proliferative potential and the demands of specific organ types, for example birds with long lifespans have high levels of telomerase in their bone marrow (Haussmann et al. 2007). In other species telomerase does not appear to be related to homeostasis and longevity, for example studies in laboratory mice have demonstrated that ageing is unlikely to be caused by telomere shortening alone as aged mice have long telomeres and loss of telomerase has no immediate effect on ageing (Blasco et al. 1997; Samper et al. 2001; Blasco 2005; Tomás-Loba et al. 2008; Jaskelioff et al. 2011). Unlike mammals that stop growing upon reaching adulthood, certain animals with indeterminate growth such as some fish (Charnov et al. 1991; Klapper et al. 1998a; Reznick et al. 2002), bivalve molluscs (Owen et al. 2007), sea urchins (Ebert et al. 2008), lobsters (Klapper et al. 1998b) and planarians (Tan et al. 2012) express telomerase through the life cycle.

In addition to its canonical role in telomere maintenance, TERT, the protein subunit of the telomerase enzyme has also been shown to have non-canonical functions (Smith et al. 2003; Parkinson et al. 2008; Choi et al. 2008; Low and Tergaonkar 2013). Not much is known about whether these functions are conserved or lineage specific. Additionally, *TERT* transcripts in many animal species, including vertebrates, insects, nematodes and planarians are alternatively spliced (Ulaner et al. 2000; Meier et al. 2006; Sýkorová and Fajkus 2009; Tan et al. 2012; Nehyba et al. 2012; Withers et al. 2012) and spliced variants have also been shown to have non-canonical roles, for example in cell proliferation (Hrdličková et al. 2012a; Listerman et al. 2013). *TERT* splicing dysregulation is present in many cancers and it is thought that human *TERT (hTERT)* splicing could be manipulated for therapeutic purposes (Wong et al. 2013; Wong et al. 2014). While only a little is known about *TERT* alternatively spliced (AS) variants in animals this represents a potential evolutionary mode for lineage specific life history adaptation, and thus broader assessment of *TERT* alternate splicing is required particularly in lineages spanning short evolutionary time frames.

Given that telomerase function is very likely to show adaptation to lineage specific life history traits, it is surprising that an in depth understanding of the evolutionary history of TERT is currently lacking from the literature. Here, we make a detailed study of TERT across the Animal Kingdom using extant data to look for evidence (or lack) of patterns of TERT protein and alternative splicing evolution that would support lineage specific adaptations. We identify TERTs across animal phylogeny and perform within and between phyla comparison of the structure of this gene. We are able to confirm the domain content of the ancestral TERT at the base of the Animal Kingdom and then we are able to show dynamic evolution of the protein in different lineages. This includes both the loss of canonical motifs and the invention of lineage specific domains, for example the invention of novel C-terminal extension and N-terminal linker motifs specific to the vertebrate lineage. In some cases we hypothesized that novel motifs may simply compensate for lost canonical motifs, and in others they may relate to lineage specific activities and interactions that remain to be discovered. Next we use available data to assess the levels of alternative splicing, implicated in non-canonical TERT function in humans and described these in different taxa. We find evidence that alternative splicing is likely to be common feature of some lineages, but is not broadly conserved with respect to splicing patterns. Finally, we study the evolution of splicing in one particular related group of animals, the free-living Tricladida planarians (Riutort et al. 1993; Carranza et al. 1996; Álvarez-Presas et al. 2008). We find that it evolves relatively rapidly in this group of highly regenerative animals, suggesting that study of alternative splicing over shorter evolutionary timescales may be required to understand adaptive non-canonical roles of TERT. Taken together our work highlights a previously unappreciated evolutionary and likely functional diversity in this core eukaryotic protein. We conclude that telomere biology, a core synapomorphy of eukaryotes, has undergone significant adaptation in different lineages and some of this is through the evolution of the TERT protein subunit of telomerase itself.

## Results and Discussion

### TERT distribution across the Animal Kingdom and reconstruction of the ancestral TERT domain structure

While TERT proteins perform core functions across eukaryotes, not much is known about how conserved or not TERT protein structure is between different metazoan lineages. In order to address this, we set out to define the structure of TERT in the metazoan ancestor and use this as a basis for comparison amongst animals. We searched for TERT orthologs from publicly available genomic and transcriptomic datasets and retrieved 135 *TERT* sequences representative of different metazoans and unicellular eukaryotes. (supplementary table S1). A maximum-likelihood tree constructed from multiple sequence alignment of TERT proteins revealed that the overall topology reflects the generally accepted phylogenetic relationship among phyla (fig. 1; supplementary fig. S1).

**Fig. 1.**
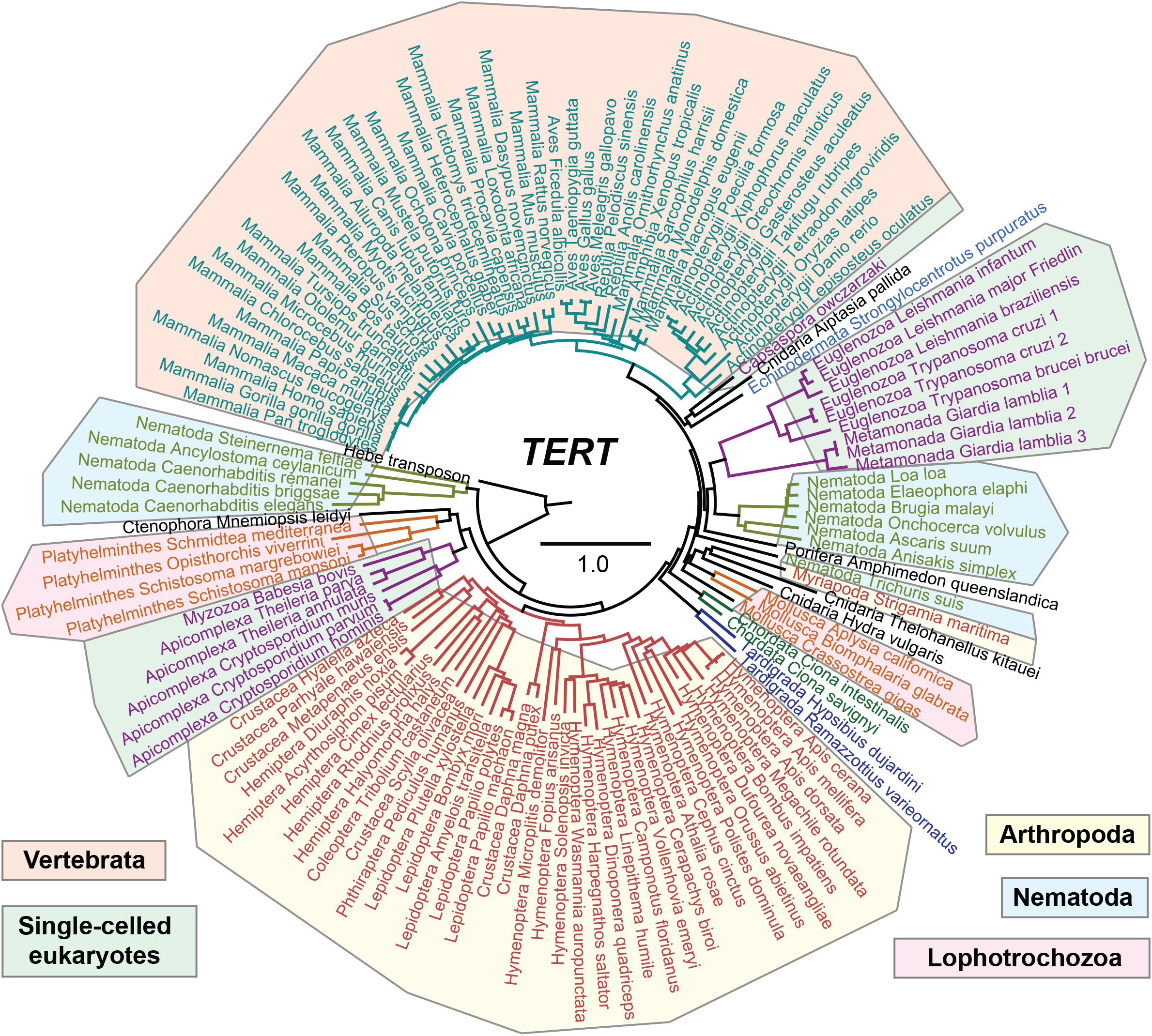
Evolution of TERT across the Animal kingdom. Phylogenetic tree of TERT sequences from representative metazoan phyla, early-branching metazoans and unicellular relatives of metazoans constructed using the maximum-likelihood method. The *Hebe* non-LTR retrotransposon is used as an outgroup. Bootstrap support values can be obtained from supplementary figure 1. Scale bar denotes substitutions per site. Full list of species used and accession numbers can be obtained from supplementary table 1.

Metazoans are related to unicellular lineages from the holozoan clade: filastereans, ichthyosporeans and choanoflagellates, (Lang et al. 2002; Ruiz-Trillo et al. 2008; Shalchian-Tabrizi et al. 2008; Torruella et al. 2012). While we could identify *TERT* from *Capsaspora owczarzaki*, we could not find evidence for *TERT* orthologs in either the choanoflagellates *Monosiga brevicollis* or *Salpingoeca rosetta*. Despite this, both *M. brevicollis* (Robertson 2009) and *S. rosetta* (Fairclough et al. 2013) have the 5’-TTAGGG(n)-3’ telomeric repeat. Based on this finding, a canonical TERT is predicted to be present (Robertson 2009). The observed absence could reflect incomplete genome sequences or, more likely given the absence from two different choanoflagellate genomes, that TERT primary sequence has evolved beyond detection or a related reverse transcriptase activity has instead taken up the role of TERT. From the genome sequences of other unicellular parasitic protozoans, putative *TERT* sequences have been predicted (Xu et al. 2004; Figueiredo et al. 2005; Hall et al. 2005; Pain et al. 2005; Dreesen et al. 2005; El-Sayed et al. 2005; Giardini et al. 2006; Morrison et al. 2007) and our phylogenetic analyses confirms that these predictions are all canonical *TERT* proteins, so it seems likely that loss of a recognisable TERT in choanoflagellates is a derived character rather than the alternative hypothesis of independent evolution of TERT like proteins in different eukaryotic lineages (fig. 1; supplementary fig. S1). *TERT* sequences from early-branching metazoans reported to date were from the genomes of *Hydra magnipapillata* and *Nematostella vectensis* (Chapman et al., 2010; Steele et al., 2011). The genomes of the sponge *Amphimedon queenslandica* (Srivastava et al. 2010), comb jelly *Mnemiopsis leidyi* (Ryan et al. 2013), myxosporean *Thelohanellus kitauei* (Yang et al. 2014) and sea anemone *Aiptasia pallida* (Baumgarten et al. 2015) have been sequenced and from these genomes we identified new *TERT* sequences for comparative analysis (fig. 1; supplementary fig. S1; supplementary table S1). While TERT proteins have been identified in many bilaterian phyla (Harrington et al. 1997; Kilian et al. 1997; Lingner et al. 1997; Nakamura et al. 1997; Meyerson et al. 1997; Bryan et al. 1998; Collins et al. 1998; Greenberg et al. 1998; Martin-Rivera et al. 1998; O’Reilly et al. 1999; Meier et al. 2006; Lau et al. 2007; Li et al. 2007; Cai et al. 2011; Kim et al. 2011; Wurm et al. 2011; Tan et al. 2012; Withers et al. 2012; Hrdličková et al. 2012b; Schumpert et al. 2015) we additionally identified novel TERT sequences that have not been reported elsewhere. These include TERT orthologs from nematodes species of the classes Secernentea, Chromadorea, Enoplea and Rhabditea and multiple orthologs from Insecta, Actinopterygii, and Mammalia (supplementary table S1). We also identified novel *TERT* sequences from parasitic platyhelminthes from the Trematoda class, from the newly sequenced tardigrade genomes of *Hypsibius dujardini* (Koutsovoulos et al. 2016) and *Ramazzottius varieornatus* (Hashimoto et al. 2016) and the crustacean *Parhyale hawaiensis* (Kao et al. 2016) (supplementary table S1).

Despite the ubiquity of TERT amongst metazoans, its presence is not universal. TERT has been reported to be absent from some animal species; one prominent example being the loss of TERT in Diptera before divergence from Siphonaptera (Mason et al. 2016). Like in dipterans, we failed to identify *TERT* in the bdelloid rotifer *Adineta vaga*, suggesting the loss of TERT from this group of long-term asexual species (Welch and Meselson 2000; Gladyshev and Arkhipova 2010a). In the absence of TERT and canonical telomere repeats, *Drosophila melanogaster* uses a retrotransposon-based telomere maintenance in which telomere-specific retrotransposons are reverse transcribed onto chromosome ends (Pardue and DeBaryshe 2003). Three non-Long Terminal Repeat (LTR) retrotransposons, *HeT-A* (Biessmann et al. 1990; Biessmann et al. 1992), *TART* (Casacuberta and Pardue 2002; Casacuberta and Pardue 2003) and *TAHRE* (Abad et al. 2004), are specifically used for telomere maintenance in this species. The bdelloid chromosome ends also have specialised telomere-associated retrotransposons, such as the *Hebe* transposon (Gladyshev and Arkhipova 2010b) and sub-telomeric reverse transcriptase (RT)-related genes (Gladyshev and Arkhipova 2011). *Penelope*-like elements, which lack the endonuclease domain and are located in sub-telomeric regions and are found in two bdelloid rotifer species *A. vaga* and *Philodina roseola* (Gladyshev and Arkhipova 2007). Although no evidence yet directly links non-LTR transposons to end chromosome maintenance in *A. vaga*, it seems plausible that in the absence of *TERT*, bdelloid rotifers may employ retrotransposon-based mechanisms for telomere maintenance.

The TERT protein contains three canonical domains and 11 motifs: 1) a telomerase essential N-terminal (TEN) domain with the GQ motif, 2) a telomerase RNA-binding (TRBD) domain with the CP, QFP and T motifs and 3) a reverse transcriptase (RT) domain with the 1, 2, A, B’, C, D and E motifs (Kelleher et al. 2002; Autexier and Lue 2006). Beyond these canonical domains and motifs, there is a flexible linker region called the N-terminal linker between the GQ and CP motifs and a C-terminal extension region that starts after the last E motif (Autexier and Lue 2006). The T motif allows for correct distinction of TERT family proteins from other prototypical RT proteins and we find that this motif is present in all species used in this study (fig. 2). To confirm the ancestral metazoan structure of TERT, we performed a cross-species comparison of all TERT proteins (fig. 2). We inferred that TERT in the metazoan ancestor possesses all 11 canonical motifs outlined above based on the observation that a complete complement of canonical motifs is present in the unicellular filasterean *C. owczarzaki* and the cnidarian *Hydra vulgaris* (fig. 2). Within Bilateria taxa considered here, only vertebrates and the echinoderm *Strongylocentrotus purpuratus* have retained all 11 motifs (fig. 2), while all protostomes have lost one or more canonical motifs suggesting broad differences in how TERT executes its core function may exist in metazoans.

**Fig. 2.**
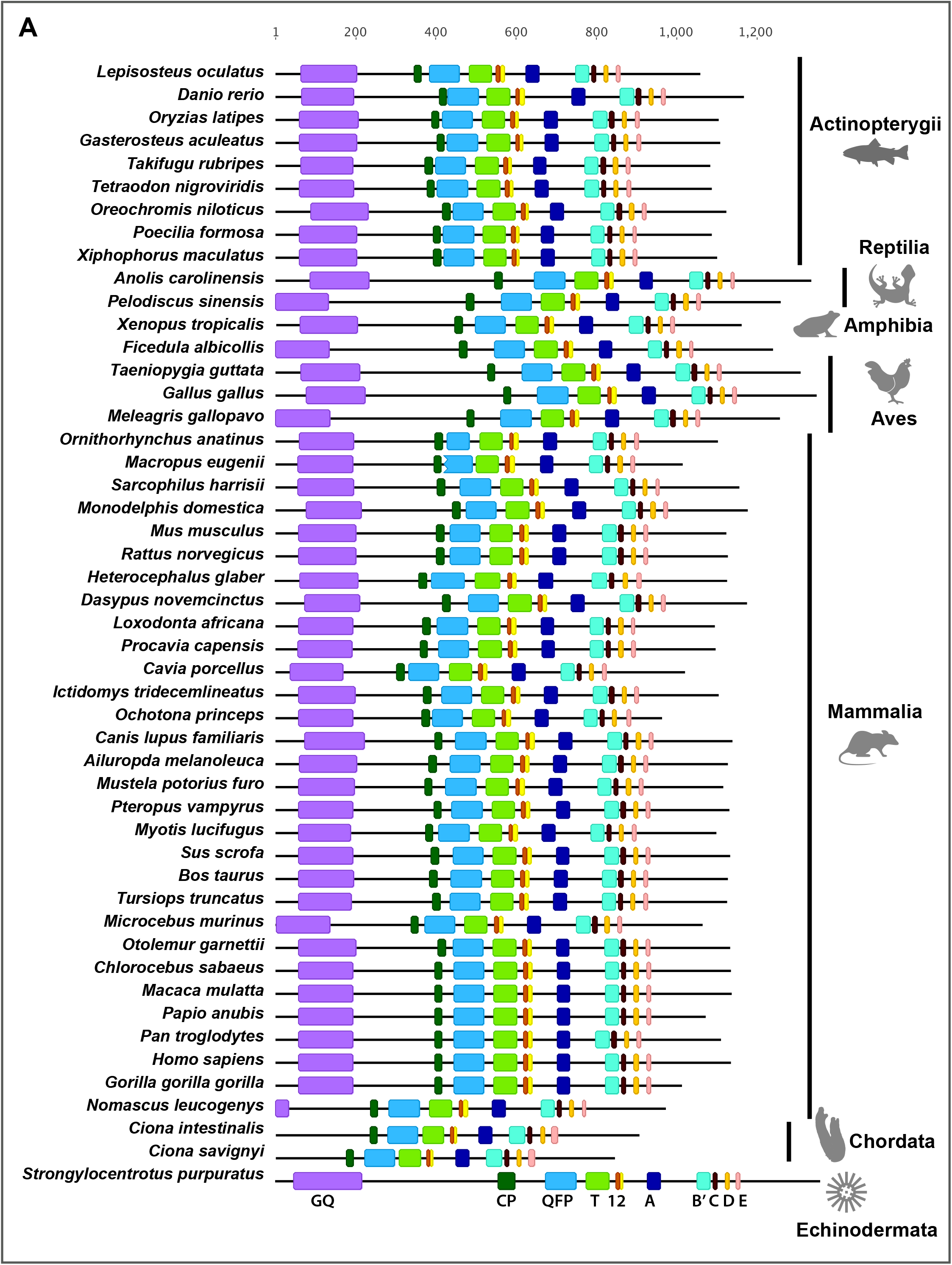
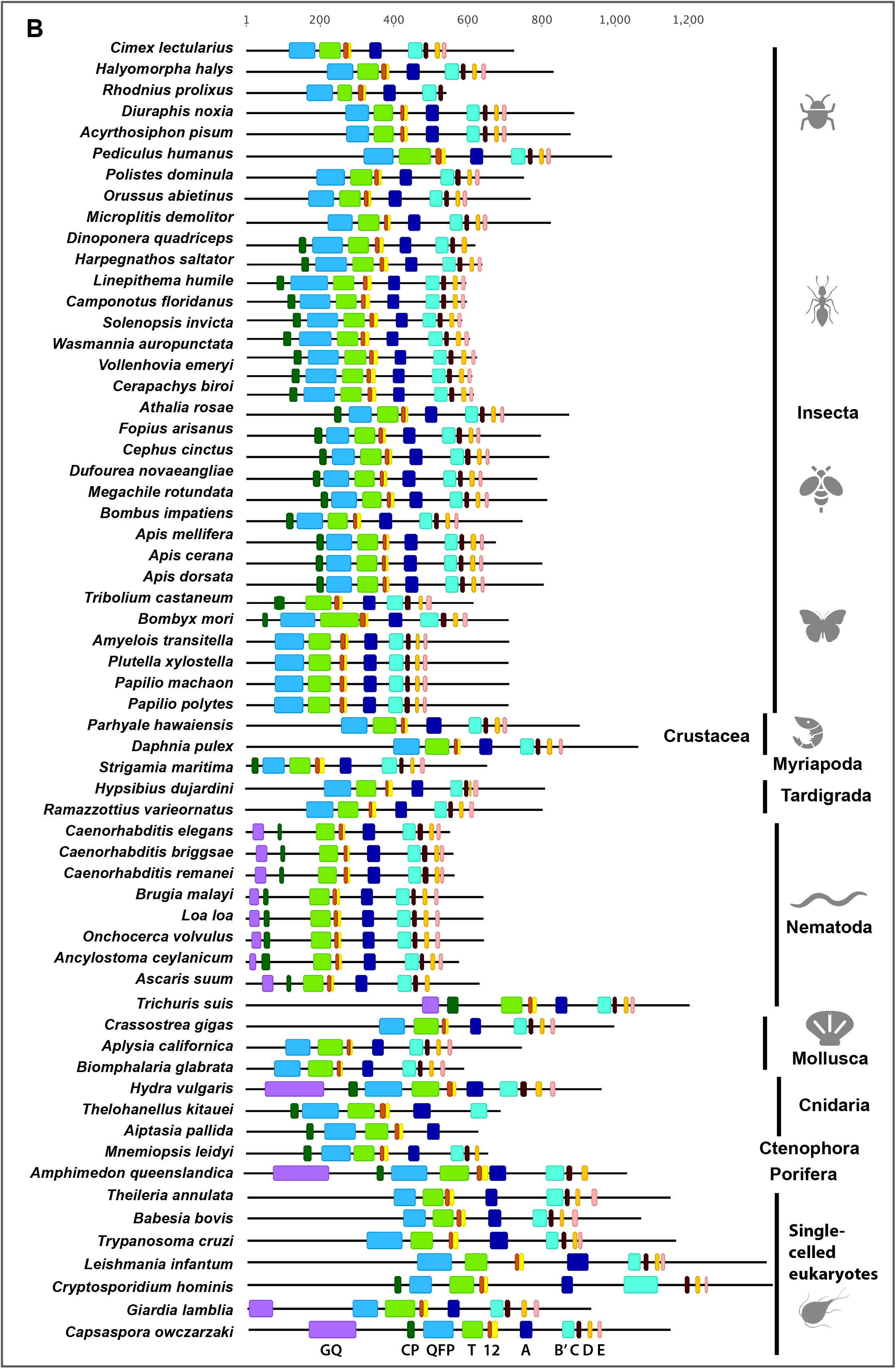
Taxa-specific modifications of TERT domains. TERT canonical motifs from representative metazoan phyla, early-branching metazoans and unicellular relatives of metazoans were annotated in colour codes: TEN domain GQ motif (purple); TRBD domain CP (dark green), QFP (blue) and T (light green) motifs; reverse transcriptase domain motifs 1 (brown), 2 (yellow), A (dark blue), B’ (sky blue), C (black), D (orange) and E (pink). **(A)** Deuterostome TERT canonical motifs. **(B)** Protostome, early-branching metazoans and unicellular eukaryotes canonical motifs. Motif annotations were drawn according to scale based on multiple sequence alignments performed using MAFFT.

### Taxa-specific divergence of canonical TERT motifs

We next examined evolution of TERT canonical motifs in detail across taxa. Sequence analyses of 11 TERT canonical motifs revealed that the TEN (GQ motif) and TRBD (CP, QFP and T motifs) domains have high sequence conservation across their whole lengths (supplementary fig. S2). Amongst unicellular eukaryotes considered here, only *C. owczarzaki* TERT has all 11 motifs (fig. 2). An independent study reported that TERT in *Leishmania amazonensis* also contained all 11 motifs (Giardini et al. 2006). However, from our sequence alignment analyses, we were unable to identify clearly defined GQ motifs in *Leishmania sp., Trypanosoma sp., Theileria sp., Babesia sp*. and *Cryptosporidium sp*. (supplementary fig. S3). The observation of GQ in *L. amazonensis* might be an alignment artefact because from their alignment, only 8 out of 244 amino acid residues are found to be in common with other TERTs (Giardini et al. 2006). We also failed to find the CP motif in this group of unicellular eukaryotes, except for *C. owczarzaki* and *Cryptosporidium* sp. (supplementary fig. S3). It was reported *Giardia lamblia* lacks the T motif (Malik et al. 2000). However, this could be due to the use of evolutionarily distant TERT sequences from plants, *Caenorhabditis elegans* and yeast, for the alignments in their earlier study. We retrieved three TERT sequences for the parasitic protozoan *G. lamblia*, obtained from different animal isolates (Adam et al. 2013). Sequence alignment revealed that they share only 72% identity. From an alignment of these *G. lamblia* TERTs with other protozoans, we were able to identify a putative T motif, suggesting that it hasn’t been lost. The *G. lamblia* T motif, however, has 8 extra residues not present in other unicellular eukaryotes (supplementary fig. S3). Apart from *C. owczarzaki* and *G. lamblia*, it appears that the TEN domain has diverged substantially in unicellular eukaryotes with poor conservation to the canonical GQ motif. Within kinetoplastid protozoans from the genera *Leishmania* and *Trypanosoma*, we identified a conserved block of residues, (I/V)QQRVxLQF, between the QFP and T motif. Considering that the divergence time of *Trypanosoma* sp. from other trypanosomatids including *Leishmania* sp. is well over 300 million years ago (m.y.a; Stevens et al. 2001), it is likely this block of residues in TERT is an example of a linage specific motif.

Amongst early branching metazoans, TERT sequences from the comb jelly *M. leidyi* and two cnidarians, *A. pallida* and the myxozoan *T. kitauei* appear to be incomplete because their sequences do not extend beyond reverse canonical motifs D, B and A respectively (fig. 2). TERT from the sponge *A. queenslandica* appears to be complete but entirely lacks the E motif. The GQ motif is collectively absent from the comb jelly and cnidarians *T. kitauei* and A. *pallida*, but not from *H. vulgaris*, indicating that multiple independent losses of GQ occurred amongst early branching metazoans (fig. 2). Within Deuterostomia, vertebrates and *Strongylocentrotus purpuratus* have retained all 11 conserved motifs while *Ciona sp*. lack the GQ motif (Li et al., 2007; fig. 2). The canonical GQ motif has also been lost from protostomes, except from the nematode lineage (fig. 2, supplementary fig. S4). In addition to the free-living *Caenorhabditis* sp. (Meier et al. 2006), we find that other nematodes have also retained the GQ motif (supplementary fig. S4). As previously reported we find that the early branching nematode *Trichinella spiralis* TERT has all 11 canonical TERT motifs (Cai et al., 2011), so the ancestor of nematode likely possessed all canonical motifs. Studies on human cell lines have shown that within the GQ motif, an additional region known as ‘dissociates activities of telomerase’ (DAT), is essential for TERT activity (Armbruster et al. 2001). Although nematodes have retained the GQ motif, the DAT region within this motif appears to have lost multiple important residues (supplementary fig. S4). The DAT region was proposed to be involved in other aspects of in vivo telomere elongation because mutations in this region have no direct effect on human TERT multimerization, nuclear targeting or template binding (Armbruster et al. 2001). It appears that either nematodes do not have *sensu stricto* DAT regions, or that the GQ motif in protostomes, including nematodes, have diverged significantly that sequence conservation with the deuterostome GQ can no longer be detected. (supplementary fig. S4). The GQ motif in yeast, human cell lines and *T. thermophila* plays a role in repeat addition processivity (Armbruster et al. 2001; Moriarty et al. 2004; Autexier and Lue 2006; Sekaran et al. 2010; Wyatt et al. 2010). It remains to be determined, however, whether the loss or sequence divergence of GQ in protostomes bears any functional implications in species-specific life histories. From the crystal structure of the red flour beetle *Tribolium castaneum* TERT, it was shown that although GQ is absent in this species, remaining motifs (CP and T) in the N-terminal region of TERT not only share high degrees of structural conservation with *T. thermophila* TERT (Gillis et al. 2008; Rouda and Skordalakes 2007) but also conferred in vitro activity (Mitchell et al., 2010). Therefore, even in the absence of a canonical GQ sequence across protostomes (fig. 2), structural and functional conservation of TERT N-terminal region may persist in these phylogenetic groups. The second TERT canonical motif CP has also been also lost in some protostomes: e.g. in molluscs and tardigrades (fig. 2). Similarly, most arthropods lack the CP motif, but several exceptions exist: the centipede *Strigamia maritima*, hymenopterans (ants, sawflies, bees but not wasps) and *T. castaneum* (Gillis et al. 2008) have retained the CP motif (fig. 2; supplementary fig. S5). Other TERT motifs are maintained in metazoan taxa.

### Identification of novel lineage-specific motifs in TERT

TERT proteins from most organisms possess two highly variable regions that are not part of the 11 canonical motifs: the C-terminal extension (CTE) and the N-terminal linker region (Autexier and Lue 2006). The CTE region starts after the last canonical E motif, and despite poor sequence homology of CTEs between human, *T. castaneum* and *T. thermophila*, it is thought to constitute the RT ‘thumb’ that is made up of three α-helices crucial in template binding and TERT processivity (Huang et al. 1998; Hossain et al. 2002; Huard et al. 2003; Gillis et al. 2008). The mechanistic significance of the CTE has remained elusive until recently, when the CTE was shown to be directly involved in differential binding of DNA and a mutated version of CTE resulted in defective DNA binding and faster DNA dissociation rates (Tomlinson et al. 2016). Since the CTE appears to play an important role despite having poor sequence conservation we examined whether this region has evolved to harbour lineage specific amino acid conservation that could potentially contribute to adaptations in different lineages. We extracted regions after the last canonical E motif from 96 species representing various taxa and investigated these as potential CTEs. Not all species have CTE regions because TERT proteins in some animals do no extend beyond the E motif. We show that unicellular eukaryotes and two early branching metazoan species, *H. vulgaris* and *A. queenslandica* possess CTE regions (supplementary fig. S6A). Alignment of unicellular eukaryotes CTE regions revealed that blocks of conservation exist within different protozoan lineages, although not between unicellular species and human TERT (supplementary fig. S6C). *Crytosporidium sp*. CTE regions are considerably shorter than those of other protozoans. Kinetoplastid protozoans, *Leishmania sp*. and *Trypanosoma sp*., have a block of conserved residues made up of 48 amino acids at the end of their TERT proteins (supplementary fig. S6C). In other early-branching metazoans, although the parasitic cnidarian *T. kitauei* and the comb jelly *M. leidyi* lack CTE regions, we identified some conserved residues in CTE regions when aligning these regions from *A. queenslandica* and *H.vulgaris* (supplementary fig. S6A).

Within bilaterians, it has been reported that nematodes from the *Caenorhabditis* genus have some of the shortest TERT proteins and appear to have lost their CTE structure (Malik et al. 2000; Meier et al. 2006). We observe that this is also true for the parasitic roundworm *Ancylostoma ceylanicum* where the C-terminal region beyond the E motif only has 42 amino acid residues (supplementary fig. S6E). We discovered that most parasitic nematodes from the Chromadorea and Secernentea classes possess intact CTE regions (supplementary fig. S6E). In addition, we show that filarial nematodes *B. malayi, O. volvulus* and *L. loa* share a RIAVL(R/K)FLKASLLEKYR(M/V motif (supplementary fig. S6E). The observation that a CTE is present in most parasitic nematodes but not the free-living *Caenorhabditis sp*. may reflect adaptation of TERT to their very different life histories; i.e. some parasitic nematodes have long generation times and life expectancies and are able to survive for years in their hosts (Morand 1996). CTE structures in Arthropoda varied markedly and we performed separate alignments for insects and crustaceans (supplementary figure S6D & F). We identified two crustacean specific motifs within the CTE region RL(K/Q)x(I/V) and R(L/F)xAL (supplementary fig. S6F). Within insects, ants lack the CTE completely as TERT proteins within this lineage do not extend beyond the E motif (supplementary fig. S6D). Lepidopterans have additional residues at the extreme ends of their TERT proteins conserved within this lineage only; a lepidopteran-specific CTE contains 32 to 36 residues (supplementary fig. S6D). Within Deuterostomia, we annotated highly conserved CTE motifs in tetrapods and fishes (supplementary fig. S6G).

Our findings suggest that the ancestor of metazoans possessed a CTE and that while this feature has been largely retained in most animal lineages, it has been lost multiple times. We have demonstrated that these CTE regions can be incredibly diverse across animal phyla, and we show that novel motifs exist in different lineages. It seems likely that these lineage specific motifs in the CTE region would have significant roles, since functional studies in *T. thermophila*, yeast and humans have revealed the importance of CTE for TERT catalytic activity related to enzyme multimerization, processivity and DNA binding (Friedman and Cech 1999; Beattie et al. 2001; Lai et al. 2001; Huard et al. 2003; Tomlinson et al. 2016). Overall investigation of the CTE supports the hypothesis that canonical TERT protein has undergone lineage specific adaptations.

We next investigated the N-terminal hypermutable linker (hereafter referred to as the linker) of TERT that is defined as the region between the GQ motif and CP motif. The linker region, although evolutionarily divergent, has been shown to be biologically essential for the function of TERT. In vertebrates, the association between TERT and telomerase RNA (TR) is mediated by the VSR motif in the linker region that binds the activation domain of TR (Moriarty et al. 2002; Bley et al. 2011). The puffer fish *Takifugu rubripes* has an additional short alpha-helix upstream of the VSR motif, known as the TFLY motif that is also implicated in binding the template boundary element (Rouda and Skordalakes 2007; Harkisheimer et al. 2013), which is a precise region within the RNA template that is defined by structural elements identified in *T. thermophila, Kluyveromyces lactis* and mammals (Autexier and Greider 1995; Tzfati et al. 2000; Lai et al. 2002; Chen and Greider 2003). Amongst early-branching metazoans, we found pockets of conservation in the linker region; the motif LxxAIF is present in three out of five early-branching metazoan species (supplementary fig. S7A). Unicellular eukaryotes have very divergent N-terminal linkers between phyla but some conservation within each protozoan lineage, the kinetoplastids (*Leishmania* sp. and *Trypanosoma* sp.) have the ALxR(T/L)(D/N)V(P/S)RxxL motif and the piroplasmids (*Babesia* sp. and *Theileria* sp.) have the LLxGLF motif (supplementary fig. S7C).

We performed an alignment of the linker regions from representative deuterostome phyla and observed that linker regions have poor conservation and they do not align well. Separately, an alignment of linkers from vertebrates alone revealed that vertebrates share a well-conserved block of 91 amino acid residues directly upstream of the CP motif, which we named the Vertebrata-specific N-terminal linker (VL; supplementary fig. S7F). Within this VL region we noticed that class specific motifs exist. According to the molecular timescale for vertebrate evolution, mammals diverged from birds 310 m.y.a. and marsupials from placental mammals at 173 m.y.a. (Kumar and Hedges 1998). Teleosts and tetrapods diverged approximately 450 m.y.a. after the vertebrate genome duplication (Hedges 2002). Such long evolutionary distances enabled us to distinguish conserved motifs from neutrally evolving sequences. For example, we demonstrated that ray-finned fishes have the FIRTLGFLY and RRxQGxD motif, birds have the NQSL motif and marsupials have the HLFxxKGDPxQQ motif (supplementary fig. S7F). It seems highly likely that these conserved motifs have functional roles that will be important for TERT function in each of these lineages.

Extending our analyses to protostomes, we observed that protostomes generally have truncated N-terminal regions. Because arthropods and lophotrochozoans lack the GQ motif altogether, we extracted sequences upstream of the CP motif (for species that have CP motifs) or the QFP motif (for species without the CP motif) as putative linker regions (supplementary fig. S7B, D & E). Insects, including ants, bees and lepidopterans exhibit long stretches of conserved residues (supplementary fig. S7D). Crustaceans from the classes Malacostraca and Branchiopoda share both FPxxHILS and ILxxNxG motifs in their N-terminal linker regions (supplementary fig. S7B). Amongst nematodes, most species with *Trichuris suris* as the exception (fig. 2), have truncated N-termini upstream of the CP motif. Nevertheless, some conservation exists within this region, for instance nematodes share a IGxxNxxxx(L/V) motif (supplementary fig. S4A). Overall, although the functional significance of the N-terminal linker is unclear beyond vertebrates, it is apparent that the linker sequence can provide additional surface areas to allow conformational flexibility between the N-terminus and the rest of the TERT protein (Autexier and Lue 2006). It is likely that the conserved linker modifications within different lineages confer some but yet unknown essential biological functions.

### TERT exhibits conservation of intron-exon structure and pan-metazoan regulation by alternative splicing

*TERT* genes are the only example of reverse transcriptase-related genes that have defined biological functions and the preservation of exon-intron structure over long evolutionary periods (Nakamura et al. 1997; Gladyshev and Arkhipova 2011). We wished to assess changes in both intron structure and alternative splicing of *TERT* across animals. We performed cross-phyla comparison of intron positions within the *TERT* gene family. We defined intron positions according to methods described by another study that compared introns across large evolutionary distances (Raible et al. 2005). Intron positions were annotated on protein sequences, where data were available, by determining the boundaries of each coding exon. TERT protein sequences with annotated intron positions were aligned and conserved intron positions were defined as shared intron placements at equivalent positions with hTERT that has 15 introns. With the exceptions of *C. owczarzaki* and *Theileria* sp., *TERT* in unicellular eukaryotes were found to lack introns altogether (fig. 3). This may reflect the fact that *TERT* is constitutively on in these unicellular organisms and is not regulated at the level of splicing. At the base of Eumetazoa, *TERT* from the comb jelly *M. leidyi* and cnidarian *T. kitauei* are also intronless while *H. vulgaris* and *A. queenslandica* have 7 and 29 introns respectively. All 7 introns in *H. vulgaris* have shared positions with *hTERT* while 14 out of 29 intron positions in *A. queenslandica* are shared with *hTERT*. These data suggest great flexibility in *TERT* intron content early in metazoan evolution. (fig. 3). Within deuterostome taxa, we observed that introns in 41 out of 48 species share equivalent positions with all 15 *hTERT* introns (fig. 3A). Many deuterostome *TERTs* have additional introns in positions not conserved with *hTERT*. For example, marsupials have between one to six additional introns in non-conserved positions and the sea urchin *Strongylocentrotus purpuratus* has 11 additional non-conserved introns (fig. 3A). We observed that *TERTs* in protostomes generally have lower intron numbers (fig. 3B). Nematodes have 9 to 14 introns and they share 7 to 10 intron positions with *hTERT* (fig. 3B). *TERT* in crustaceans are surprisingly intron poor, despite reports that the genome of *Daphnia pulex* has one of the highest intron densities (Colbourne et al. 2011). *D. pulex* and *P. hawaiensis* have only 1 and 3 introns respectively, and all introns have shared positions with *hTERT* (fig. 3B). *TERT* genes in the Hemiptera suborder Sternorrhyncha have undergone unique genomic changes compared to other hemipteran insects; *TERT* in aphids and lepidopterans have lost all introns (fig. 3B). Within lophotrochozoa, only the pacific oyster is intron rich; the oyster has 27 introns and 14 of these share equivalent positions with *hTERT* (fig. 3B). Other molluscs are intron poor; *Aplysia californica* and *Biomphalaria glabrata* have 4 and 3 introns respectively, none of which have positions shared with *hTERT* (fig. 3B). Introns provide the opportunity for creating new gene products via alternative splicing mechanisms. *TERT* alternatively spliced products have been implicated in non-canonical functions in human disease processes, particularly cancer and in adaptation to reproductive mode in planarians flatworms. Alternative splicing has been reported in many other taxa (Ulaner et al. 1998; Ulaner et al. 2000; Meier et al. 2006; Sæbøe-Larssen et al. 2006; Chang and Delany 2006; Lau et al. 2008; Imamura et al. 2008; Sýkorová and Fajkus 2009; Wurm et al. 2011; Hrdličková et al. 2012b; Nehyba et al. 2012). We sought to determine whether alternative splicing of TERT is common across animals and if AS variants show any conservation. We probed available genomic and expression datasets for the presence of AS variants. We identified 19 animal species with *TERT* AS variants as evident by computational predictions using RNA-sequencing datasets (supplementary table S2). We found AS variants in both protostomes (10 species) and deuterostomes (9 species). From this list, AS variants have been previously described for these five species: *Homo sapiens* (Ulaner et al. 1998; Ulaner et al. 2000; Sæbøe-Larssen et al. 2006; Hrdličková et al. 2012a), *Ornithorhynchus anatinus* (Hrdličková et al. 2012b), *Gallus gallus* (Chang and Delany 2006; Nehyba et al. 2012), *Danio rerio* (Lau et al. 2008; Imamura et al. 2008) and *Solenopsis Invicta* (Wurm et al. 2011). We observed extensive patterns of alternative splicing in hymenopteran and hemipteran insects (supplementary fig. S8). Some of the most common splicing events in hymenopterans and hemipterans involved the exclusion of exon 2 (6 out of 9 species) and exon 3 (4 out of 9 species; supplementary fig. S8). All 43 AS variants in insects have retained an open reading frame (ORF; supplementary fig. S9). Of these, 11 have truncated TRBD (motifs CP, QFP and T) domains and 9 have truncated RT (motifs 1, 2, A, B’, C, D and E) domains (supplementary fig. S8). The substantial lengths and complete ORFs of many AS variants in insects indicate that they may still be functional at the protein level. In Lophotrochozoa, two AS variants were identified in the mollusc *A. californica* (supplementary fig. S9). Since the combination of exons 2 and 3 only encode 39 amino acid residues, all AS variants in *A. californica* appear to retain a complete set of motifs like the full-length TERT (supplementary fig. S9). Within Arthropoda, aphids and lepidopterans lack any introns and hence alternative splicing (fig. 3). The short-lived nature of these taxa may preclude the need for highly regulated TERT activity.

**Fig. 3.**
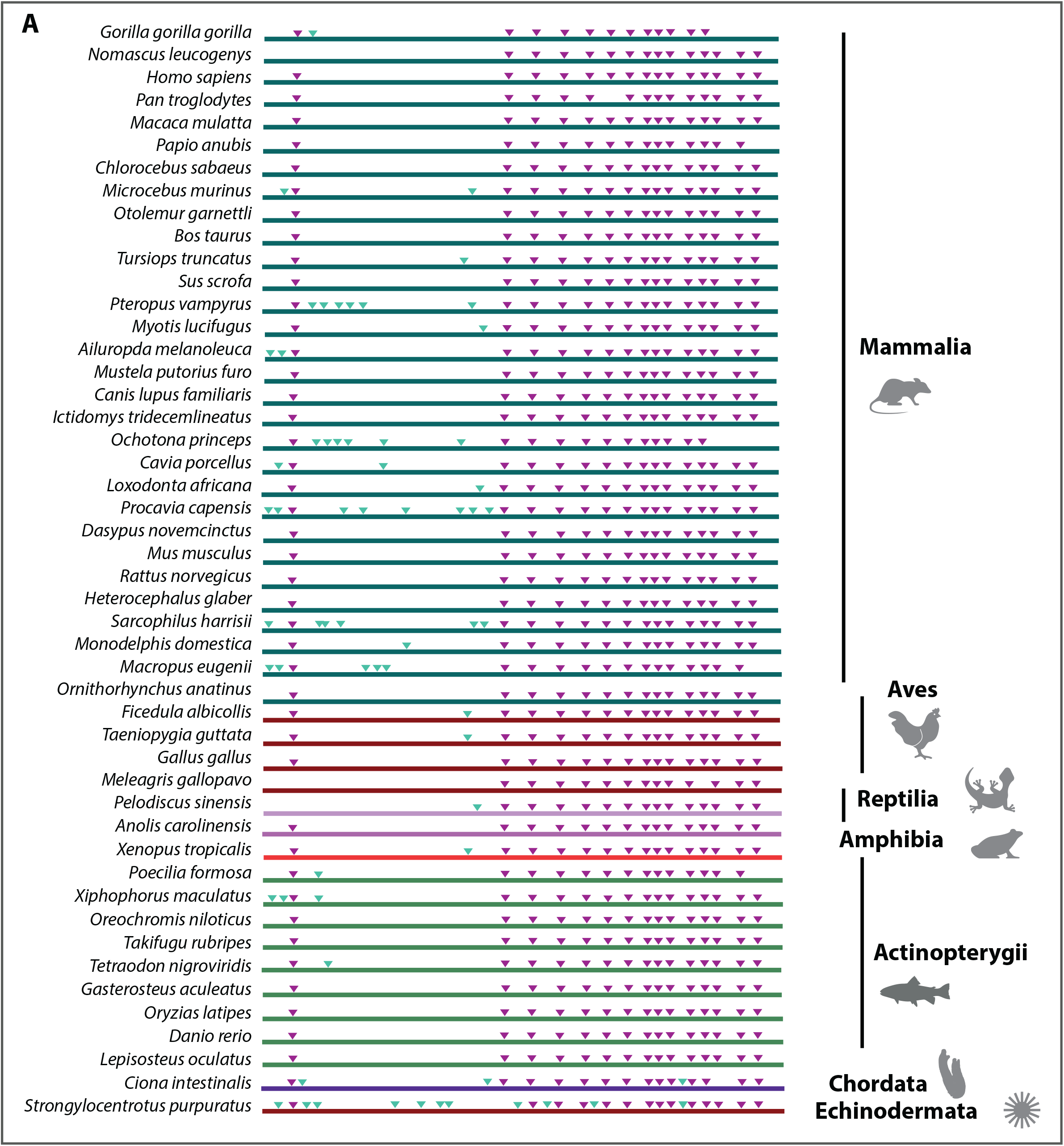
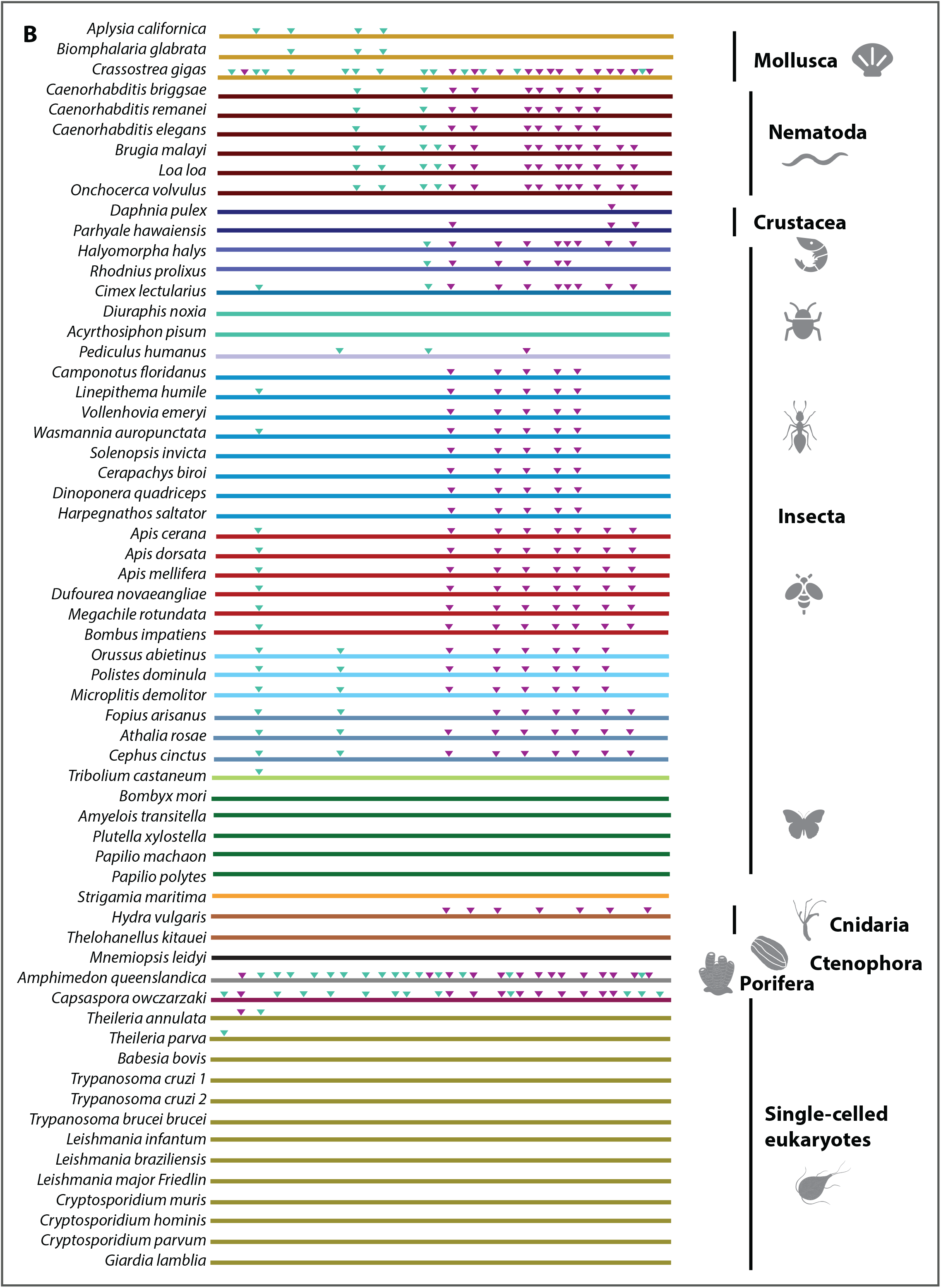
*TERT* exon-intron structure is conserved across the Animal kingdom. Schematic diagram depicting the intron positions mapped on the TERT orthologs relative to intron positions in human TERT (hTERT). **(A)** Deuterostome *TERT* intron positions. **(B)** Protostome, early-branching metazoans and unicellular eukaryotes intron positions. Purple triangles represent conserved introns (relative to hTERT intron positions) and green triangles represent species-specific introns.

Amongst vertebrate species, many AS variants have been previously described but to our knowledge, splicing patterns of *TERTs* in the Amazon molly *Poecilia formosa*, the green spotted puffer fish *Tetraodon nigroviridis*, the wild turkey *Meleagris gallopavo* and the Rhesus macaque *Macaca mulatta* have not been reported (supplementary fig. S10). AS variants in *hTERT* involved exons 6 to 8 and we witness the splicing events concerning these exons in five out of nine species (3 mammals, the wild turkey and zebrafish) (supplementary fig. S10), suggesting some conservation of splicing across the vertebrates. The exclusion of the unusually large exon 2 (1.3kb) is a common 5’ splice variant in primate *TERTs* (Withers et al. 2012) and we observe this splicing event in all vertebrates except the Rhesus macaque (supplementary fig. S10). The lack of observation of the AS variant in Rhesus Macaque may be due to lack of confirmatory data. Overall, many AS variants in mammals and fishes appear to be of substantial lengths, suggesting that they would encode functional proteins given that most of their TRBD and RT motifs are still present (supplementary table 2). In avians however, many AS variants appear to lack any canonical motifs. The wild turkey has 10 AS variants, 8 of these involve the exclusion of multiple exons and/or loss of frame mutations, do not contain any TERT motifs and are therefore likely to be non-coding and be subjected to nonsense-mediated decay (supplementary fig. S10B). This observation of complex splicing patterns generating many short AS variants with premature termination codon has also been reported in chickens (Chang and Delany 2006; Nehyba et al. 2012; supplementary fig. S10B). It is unknown as to whether these AS variants would participate directly or indirectly in canonical telomerase function or in other non-canonical functions of TERT. But clearly alternative splicing of the TERT protein is a pan-metazoan phenomenon. Given that we do not see broad conservation of splicing patterns (except within the vertebrate lineage) or data suggest, that in combination with the lineage specific sequence evolution we have also described, AS is likely to contribute to lineage specific functions of this core eukaryotic protein.

### Free-living planarian species exhibit complex patterns of alternative splicing

While our analysis of alternative splicing on TERT revealed extensive evidence of splicing, data available to assess evolutionary changes of splicing in taxa outside of the vertebrates is relative sparse. A previous study had investigated telomere length, telomerase activity and TERT regulation in the highly regenerative model planarian *Schmidtea mediterranea* (Tan et al. 2012). This study found that the asexually reproducing biotype upregulated active full length splice forms of *TERT* during regeneration to sufficient levels to maintain telomere length during stem cell division. Another study in the related species *Dugesia ryukyuensis* demonstrated equivalent telomere length maintenance differences between sexual and asexual strains (Tasaka et al. 2013). In order to use planarians as a study system for understanding how AS could contribute to the evolution of TERT regulation we performed a detailed investigation of *TERT* gene structure, expression and alternative splicing in Tricladida planarians, a group where most, but not all species, are known to be highly regenerative.

We obtained 5 new partial *TERT* sequences from published transcriptome sources or previous studies (Robb et al. 2008; Nishimura et al. 2012; Tan et al. 2012; Wheeler et al. 2015; Robb et al. 2015; Egger et al. 2015; Brandl et al. 2016). Based on these sequences, we designed putative pan-flatworm degenerate primers and using a combination of degenerate PCR together with 5’ and 3’ RACE techniques, we cloned full-length *TERT* sequences from 11 Tricladida planarian species. These included ten Continenticola species (a clade consisting of freshwater triclads from the family Dendrocoelidae, Dugesiidae and Planariidae) and one Maricola species (a suborder of marine triclads from the family Procerodidae; fig. 4A & B; supplementary table 3). We also confirmed both the identity of each species and phylogenetic relationships amongst taxa using 18S rDNA resequencing (supplementary fig. S11; supplementary table 3; Riutort et al. 1993; Carranza et al. 1996; Álvarez-Presas et al. 2008).

**Fig. 4.**
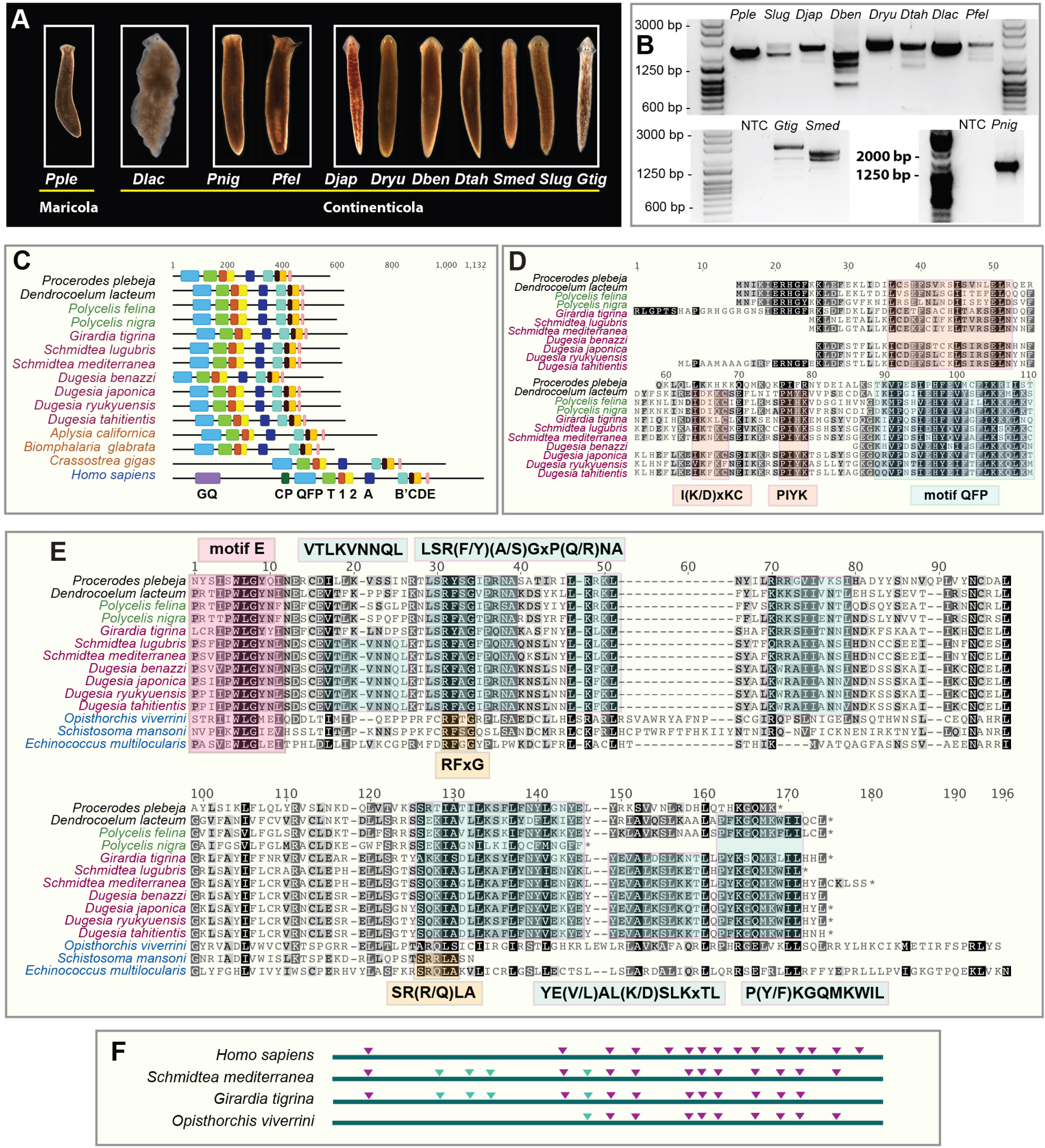
*TERT* in highly regenerative triclad planarians. **(A)** The 11 flatworm species used in this study. Abbreviations represent: *Pple (Procerodes plebeja), Dlac (Dendrocoelum lacteum), Pnig (Polycelis nigra), Pfel (Polycelis felina), Djap (Dugesia japonica), Dryu (Dugesia ryukyuensis), Dben (Dugesia Benazzi), Dtah (Dugesia tahitientis), Smed (Schmidtea mediterranea), Slug (Schmidtea lugubris)* and *Gtig (Girardia tigrina)*. **(B)** Gel image showing full length RT-PCR results of *TERT* transcripts and isoforms from 11 planarian species (NTC represents no cDNA template control for the PCR). **(C)** Domain structure of planarian TERTs; mollusc TERTs and hTERT were used for comparison. Multiple sequence alignments of the **(D)** N-terminal linker and **(E)** C-terminal extensions from planarians and the trematode*s*. Boxes also denote planarian-specific motifs with their respective descriptions written inside. **(F)** *TERT* gene structure of *hTERT, S. mediterranea, G. tigrina* and the liver fluke *Opisthorchis viverrini* (Trematoda Class) are shown. Purple triangles indicate conserved intron positions (relative to *hTERT* intron positions) and green triangles indicate Platyhelminthes-specific introns. Genomic sequence data is not available for other planarian species. Therefore, putative *TERT* exon-intron boundaries were annotated based on *Smed_TERT* data and confirmed by AS variant analyses of each species.

We observed that like other lophotrochozoans (*A. californica, Biomphalaria glabata* and *Crassostrea gigas*), TERT in triclad planarians lack the GQ and CP motifs. (fig. 4C; supplementary fig. S12). We retrieved 6 partial sequences for other platyhelminthes from the following classes: Trematoda (*Schistosoma mansoni, S. margrebowiei, Echinostoma caproni; Opisthorchis viverrini;* Young et al. 2014; Howe et al. 2016), Cestoda (*Echinococcus multilocularis;* Tsai et al. 2013) and Turbellaria (*Macrostomum lignano;* Wasik et al. 2015). Since these sequences are incomplete, we only used species with intact CTE or N-terminal linker regions for comparison. Although we failed to compare N-termini of parasitic flatworms, we were able to identify planarian-specific N-terminal motifs within this region such as the I(K/D)xKC and PIYK motifs (fig. 4D). With regards to CTE regions, there is poor conservation between molluscs and platyhelminthes and they require separate alignments. Within platyhelminthes CTE regions, we observed pockets of conservation, e.g. the P(Y/F)KGQMKWIL and LSR(F/Y)(A/S)GxP(Q/R)NA motifs in triclad planarians (fig. 4E). The VTLKVNNQL and YE(V/L)AL(K/D)SLKxTL motifs are conserved within Dugesiidae (fig. 4E).

The observation of multiple PCR products when using primers that span the entire *TERT* transcript in planarians was suggestive of *TERT* alternative splicing (fig. 4B). We wished to investigate the extent of the regulation by alternative splicing in this group of animals that exist over short evolutionary time frames. It was previously reported that *TERT* in *S. mediterranea* is alternatively spliced into 4 variants (Tan et al. 2012). Analysis of *TERT* intron-exon boundaries using existing genome assembly datasets for *S. mediterranea* (Robb et al. 2015) and *Girardia tigrina* (Kao et al. unpublished data) revealed that *Smed-TERT* and *Gtig-TERT* have 16 and 15 exons respectively. Of these, 11 *Smed-TERT* and 10 *Gtig-TERT* introns are conserved with *hTERT* introns where they share the same positions on the TERT protein (fig. 4F). As planarian *TERTs* shared a high degree of structural similarity and due to the lack of genomic data for the nine remaining planarian species, we based our AS variant analysis on an inferred genomic structure generated from the *Smed-TERT* sequence. From the splicing patterns of cloned AS variants, we could visualise exons that are being excluded and this confirmed that the intron-exon boundaries were accurately inferred based on *Smed-TERT* sequence (supplementary fig. S13 & S14). AS variants were identified from all seven Dugesiidae species, two Planariidae species and one Dendrocoelidae species (supplementary fig. S13, S14A & 14B). We did not find any AS variants from the marine planarian *Procerodes plebeja* (Procerodidae), an outgroup to all other species (supplementary fig. S14C). The highest number of AS variants was identified in *G. tigrina* (13 variants) with the lowest being *Dendrocoelum lacteum* (2 variants; Supplementary figures 13 & 14). Two of the shortest AS variants in *G. tigrina* did not contain any TERT motifs (supplementary fig. S13). Dugesiidae species in general had more AS variants compared to Dendrocoelidae and Planariidae species. Five AS variants in *D. ryukyuensis* only have partial QFP motifs and not any other TERT motifs (supplementary fig. S13). Of all 52 AS variants in triclad planarians, the most prevalent pattern is the truncation of the RT domain where 24 out of 52 have partial or missing RT domains (supplementary table 4; supplementary fig. S13 & S14). We observed 20 out of 52 AS variants containing truncated versions of both TRBD and RT domains and only 7 out of 52 contain a truncated TRBD but intact RT domain (supplementary table 4; supplementary fig. S13 & S14). Despite sharing conserved positions of introns (as inferred from splicing patterns of each species), there is low degree of conservation in the selection of spliced sites for these species. The most prevalent splicing pattern is the deletion of exons 8 to 10 (represented by orange triangles), which is present in six out of ten species where AS variants were found (supplementary fig. S13 & S14). Deletion of exons 11 and 12 (represented by blue triangles) were identified in five out of ten species. These splicing events would affect the *TERT* RT domain that is encoded by exons 8 to 13 (supplementary fig. S13 & S14). We note that the least amount of *TERT* splicing is observed in the least regenerative species, *P. plebeja* and *D. lacteum* (supplementary figure S15 & S16; Liu et al. 2013). The significance of this, if any, awaits further investigation. Overall our data suggest that AS splicing of TERT is evolving rapidly within this group of animals and that both fine grained study of *TERT* alternative splicing as well as functional study of AS variants are required to understand why *TERT* has evolved so dynamically during animal evolution.

### Conclusion

We performed thorough analyses of TERT canonical and non-canonical motifs, gene structure and alternative splicing across representative metazoan species. We show that although the ancestral TERT protein in Metazoa is likely to possess all 11 canonical motifs, the GQ and CP motifs are prone to lineage specific losses (fig. 2). Beyond the canonical motifs, we demonstrate that the N-terminal linkers and CTE regions of TERTs are highly divergent across phyla. In these regions, we discovered novel motifs that exhibit high levels of sequence conservation over long evolutionary times indicating that they may serve an unknown but important biological function. For example, the CTE regions are distinct within phyla (supplementary fig. S6) and it seems likely that such differences are reflective of life history diversity as growing evidence points to the importance of the CTE for telomerase processivity in vivo (Hossain et al., 2002; Huard et al., 2003; Lue, 2004). We also demonstrated that the N-terminal linker regions have poor sequence conservation between metazoan phyla implying that these regions are adapted to different lineages (supplementary fig. S7). The N-terminal linkers of TERT proteins have been implicated to mediate a conserved function in enhancing the translocation of the 3’ end of DNA substrate relative to the RNA template (Collins, 1999; Peng et al., 2001). Deleting residues in this region reduces the translocation of DNA substrate and overall processivity (Moriarty et al., 2004). Future studies will be needed to unravel the biological significance of phyla specific sequences and whether they fine tune telomerase processivity to life history strategy.

Analysis on *TERT* intron exon structure revealed that the metazoan ancestor is intron rich and share many intron positions with *TERTs* in the vertebrate lineage (fig. 3). *TERTs* in protostomes have undergone significant intron loss; lepidopterans and aphids do not have any introns, perhaps due to short lifespans of these taxa. Alternative splicing of TERT has been extensively studied in many species. We report new observations on splicing events for 14 metazoan species and show that the selection of spliced exons is poorly conserved in these animals (supplementary fig. S8, S9 & S10). Lastly, we investigated *TERT* alternative splicing in Tricladida planarians and demonstrated that splicing is dynamic and rapidly evolving within this group of closely related species (fig. 4, supplementary fig. S13 & S14). It is likely that splicing in planarian *TERTs* is significant for the highly regenerative and potentially immortal life histories of some of these species and AS variants could possess important non-canonical functions. This study highlights that neither *TERT* structure nor its regulation is static. A future endeavour will be to unravel mechanistic linkages that connect unique sequence evolution and the dynamic regulation of TERT to actual biological roles.

## Materials and methods

### Data mining of TERT sequences

TERT sequences were retrieved from Ensembl Genome and Ensembl Metazoa (Aken et al., 2016), NCBI and Uniprot. Sequences were confirmed using reciprocal blastx against the nr database using the NCBI Blast suite. We retrieved tardigrade TERTs from the *Hypsibius dujardini* genome browser (http://badger.bio.ed.ac.uk/H_dujardini/blast/index) and the *Ramazzottius varieornatus gene models* (http://kumamushi.org/database.html). TERT sequence from the crustacean *Parhyale hawaiensis* was retrieved from the assembled genome (Kao et al., 2016). Additional references for other datasets are included in the main text. Complete species names, accession numbers and database sources are listed in supplementary table 1.

### Phylogenetic analyses

Multiple sequence alignment of TERT protein sequences was performed using MAFFT (Katoh et al. 2009). Phylogenetic tree was constructed using RAxML (Stamatakis 2014) employing the WAG matrix and gamma distribution rate model with 1000 bootstrap replicates. A single tree topology was generated based on the best-scoring maximum likelihood tree. Alignment and tree figures were made using Geneious (version 7; Kearse et al., 2012).

### Annotation of intron positions on TERT protein sequences

For cross-species intron comparison of the core dataset, intron positions were annotated on the respective protein sequence of a given species. A scale diagram was drawn based on multiple sequence alignment of all TERT proteins containing annotated intron positions. Intron positions were defined according to previously described methods (Raible et al., 2005). Intron positions were annotated on protein sequences, where data was available, by determining the boundaries of each coding exon. TERT protein sequences with annotated intron positions were aligned and conserved intron positions were defined as shared intron placements at equivalent positions with *hTERT*. Intron positions conserved with *hTERT* were represented as purple triangles while species-specific introns were represented as green triangles. Fasta sequences of all TERT proteins were listed in supplementary file 1.

### Annotation of TERT canonical motifs and non-canonical motifs in C-terminal extension (CTE) and N-terminal linker regions

TERT protein motifs (GQ, CP, QFP, T, 1, 2, A, B’, C, D and E) were annotated based on regional comparisons with hTERT protein annotations in multiple sequence alignments performed using MAFFT (Katoh et al. 2009). Non-canonical motifs in CTE and linker regions were identified based conserved residues in multiple sequence alignments. Alignment figures were generated using Geneious (version 7; Kearse et al., 2012).

### Animal culture

All freshwater flatworm strains were cultured at 20°C using the planarian water formulation. 1X Montjuic salt solution was prepared using milliQ ddH2O with the following composition: 1.6mM NaCl, 1mM CaCl_2_, 1mM MgSO_4_, 0.1mM MgCl_2_, 0.1mM KCl, 1.2mM NaHCO_3_ (González-Estévez et al., 2012). The marine flatworm *Procerodes plebeja* was cultured at 14°C using a salt water formulation made with Tropic Marin sea salt to a salinity of 28-30ppm. *Dendrocoelum lacteum* was fed with shrimp while all other flatworms used in this study were fed organic beef liver once a week. All flatworms were starved for 1 week prior to any experimental procedures. The worms were kept in the dark at all times apart from feeding and water changing times.

### RNA extraction and reverse transcription in Triclad flatworms

Total RNA was isolated from three animals of each species using the TRIzol reagent (Thermo Fisher Scientific) according to manufacturer’s instructions. cDNA was generated from total RNA extract using the QuantiTect reverse transcription kit (Qiagen) containing the RT primer mix (blend of oligo-dT and random hexamers). A genomic DNA elimination step was also included in this kit performed using the gDNA wipeout buffer.

### Degenerate polymerase chain reaction (PCR) and 5’ 3’ RACE for the cloning of *TERT* genes

*TERT* transcript sequences for *Girardia tigrina* (Wheeler et al., 2015), *Dugesia japonica* (Nishimura et al., 2012), *D. lacteum*, and *P. nigra* (Brandl et al., 2015) were retrieved using tblastn with *Smed_TERT* as a query from published transcriptome sequences of the respective species. Top hits with the best e-value for each species was used for reciprocal blast against the nr database in NCBI to confirm the identification of *TERT*. For degenerate PCR, first strand cDNA synthesis was performed as mentioned previously. For 5’ and 3’ RACE, first strand cDNA synthesis was performed on TRIzol extracted total RNA using the SMARTer RACE 5’ 3’ Kit (Clontech).

For the cloning of *TERT* in all the other *Dugesia* sp. (*Dugesia ryukyuensis, Dugesia tahitientis, Dugesia benazzi*) degenerate PCR primers were designed based on the pairwise alignment of *Smed-TERT* and *Djap-TERT*: DugesiaFD (5’-CACTGGTGYGAATCACCA-3’) and DugesiaRD (5’-AAAACATCATCMACRTATTG-3’). PCR products were cloned in the pGEMT-easy vector (Promega) followed by colony PCR with M13F (5’- GTAAAACGACGGCCAGT-3’) and M13R (5’-GGAAACAGCTATGACCATG-3’) primers. Positive colonies were selected for Sanger sequencing. Once the middle sequence of *TERT* transcripts for all three *Dugesia* sp. were obtained, 5’ and 3’ RACE was performed on cDNA made with the SMARTer RACE 5’ 3’ Kit using gene specific primers listed in supplementary table 3 according to manufacturer’s instructions.

For the cloning of *TERT* in *Schmidtea lugubris*, the same set of degenerate PCR primers (DugesiaFD and DugesiaRD) was used for PCR followed by cloning and sequencing. PCR products were cloned in the pGEMT-easy vector (Promega) followed by colony PCR and sequencing as mentioned previously. The 5’ end of *Slug_TERT* transcript was cloned using the following primer set: Dugesia5F (5’-AATYGAGMGWMATGGTTT-3’; this primer was designed based on 5’ sequence region in *Dugesia* sp.) and LugR (5’- CTGAAATTTGTGCCATTG-3’; *Slug-TERT* gene specific primer). Next, 3’ RACE was performed with the SMARTer RACE 5’ 3’ Kit using gene specific primers listed in supplementary table 3 according to manufacturer’s instructions.

For the cloning of *TERT* in *Polycelis felina*, degenerate PCR primers were designed based on pairwise alignment of *Pnig-TERT* and *Polycelis tenuis TERT* (obtained from PlanMine; Brandl et al., 2015): PolycelisDF (5’-AATTGGCACMTSTTYCTG-3’) and PolycelisDR (5’-GACTCRTARCAAYTCTT-3’). PCR products were cloned in the pGEMT-easy vector (Promega) followed by colony PCR and sequencing as mentioned previously. The 5’ and 3’ end of *Pfel-TERT* was obtained using 5’ and 3’ RACE with the SMARTer RACE 5’ 3’ Kit using gene specific primers listed in supplementary table 3 according to manufacturer’s instructions.

For the cloning of *TERT* in *P. plebeja*, degenerate PCR primers were designed based on pairwise alignment of *Pnig_TERT* and *Pfel_TERT*: PpleDF (5’- ATGSARTATAAAGGATWTAT-3’) and PpleDR (5’-AAYRTCRTCGACATATTG-3’). PCR products were cloned in the pGEMT-easy vector (Promega) followed by colony PCR and sequencing as mentioned previously. The 5’ and 3’ end of *Pple-TERT* was obtained using 5’ and 3’ RACE with the SMARTer RACE 5’ 3’ Kit using gene specific primers listed in supplementary table 3 according to manufacturer’s instructions.

All degenerate PCRs were performed using the Advantage 2 Polymerase (Clontech) according to recommended thermal cycling parameters. All RACE products were ran on a gel and gel extraction was performed with the MinElute gel extraction kit (Qiagen) following by cloning and sequencing of the RACE products.

### DNA extraction, 18S rDNA PCR and sequencing

High molecular weight genomic DNA (gDNA) was isolated using the phenol-chloroform method followed by ethanol precipitation. PCR was performed according to conditions in Carranza et al., (1996) using the Advantage 2 Polymerase (Clontech). List of 18S primer sequences are provided in supplementary table 3. PCR products were gel extracted using the MinElute gel extraction kit (Qiagen) and eluted in 10uL of molecular grade water. Gel extracted products were sequenced in both directions using 18S nested primers listed in supplementary table 3. Sequences were aligned with known 18S sequence data available from NCBI (accession numbers provided in supplementary table 3). 18S rDNA sequences obtained from this study fell into two groups (18S type I and 18S type II).

### Cloning of planarian *TERT* AS variants

PCR primers were designed to amplify from the start to end of *TERT* transcript sequences obtained from above. Full list of primers used to amplify *TERT* AS variants are provided in supplementary table 3. PCR was performed using Phusion Polymerase (Thermo Fisher Scientific) and all PCR products were ran on a gel. In order to cut out gel pieces containing the AS variants, the gel image was overexposed to enable visualization of low abundance variants. Gel regions containing bands were excised and extracted using the MinElute gel extraction kit (Qiagen). A-tailing was subsequently performed with GoTaq Polymerase (Promega) for TA-cloning purposes since Phusion PCR products were blunt ended. PCR products were cloned in the pGEMT-easy vector (Promega) followed by colony PCR with M13F and M13R primers. Colony PCR products were run on a gel for 2 hours using 1.3% agarose in 1X TAE buffer to allow good resolution of bands. Colony PCR gel would display an array of bands with different sizes representing the AS variants. Bands of different sizes were selected for Sanger sequencing. For each species, 25 colonies were selected for sequencing to allow exhaustive identification of *TERT* AS variants.

### Sequence analyses of flatworm *TERT* AS variants

Eleven planarian species were used in this study. Planarians *Schmidtea mediterranea, S. lugubris, G. tigrina, D. japonica, D. benazzi, D. tahitientis, P. felina, P. nigra* and *D. lacteum* were purchased from Sciento (http://www.sciento.co.uk/). *D. ryukyuensis* was a gift from Dr. *Midori Matsumoto (Keio University, Japan) and P. plebeja* from Dr. Bernhard Egger (Universität *Innsbruck*). The gene structures (exon-intron boundaries) for *Smed-TERT* and *Gtig-TERT* were identified using blastn against the *S. mediterranea* (Robb et al. 2015) and *G. tigrina* genome assembly (Kao et al. unpublished data). Both *Smed-TERT* and *Gtig-TERT* have almost identical intron positions, with *Gtig-TERT* having one less exon. Since genome data is not available for all the other flatworm species, we based our analyses on the gene structure of *Smed-TERT* and *Gtig-TERT* and refer to them as inferred exon positions. Further analyses on cloned AS variants for species without genomic data confirmed the accurate positioning of intron-exon boundaries based on the excluded exons. The full-length *TERT* sequences for each species were referred to as ‘isoform 1’. Cloned AS variants were mapped to ‘isoform 1’, which set as a reference sequence for each respective species. From this, we were able to visualize the skipped exons, splice site mutations or retained introns by comparing the AS variant sequences to the reference sequence. All the AS variants including the longest isoform were *in silico* translated using Geneious (version 7; Kearse et al. 2012) to the correct open reading frame. Multiple sequence alignments of all AS protein sequences for each respective species were performed using MAFFT (Katoh et al. 2009). Annotations of TERT canonical motifs were performed according to positions in hTERT; telomerase RNA-binding domain motifs (QFP and T) and canonical reverse transcriptase motifs (1, 2, A, B’, C, D and E) in all AS protein variants. Further information on the flatworm AS variants were provided in supplementary table 4.

### Annotation of *TERT* AS variants in vertebrates, insects, nematodes and molluscs

For *TERT* sequences that contained computationally predicted isoforms generated based on evidence from RNA-sequencing contigs, we extracted the coding sequences (CDS) from these predictions and performed in silico translation according to the correct open reading frame. Subsequent analyses were performed as described in the previous section. The longest transcripts/isoforms were used as reference sequences and all other AS variants were mapped to the reference using Geneious (version 7; Kearse et al. 2012). This allowed the visualization of skipped exons, splice site mutations or retained introns by comparing the AS variant sequences to the reference sequence. Detailed descriptions on splicing patterns are available in supplementary table 2.

## Acknowledgments and funding information

This work was supported by the Biotechnology and Biological Sciences Research Council (grant number BBK0075641); Human Frontier Science Program Postdoctoral Fellowship (to A.G.L.); University of Oxford Elizabeth Hannah Jenkinson Grant (to A.G.L. and S.S.) and Clarendon Scholarship (to S.S.). We thank Dr. Hanae Nodono (Kagoshima University, Japan) for her contribution on TERT identification in planarians. We thank Dr. Midori Matsumoto (Keio University, Japan) for the gift of *Dugesia ryukyuensis*. We thank Dr. Bernhard Egger (Universität Innsbruck) for the gift of *Procerodes plebeja*.

## Figure legends

**Supplementary Figure S1.**
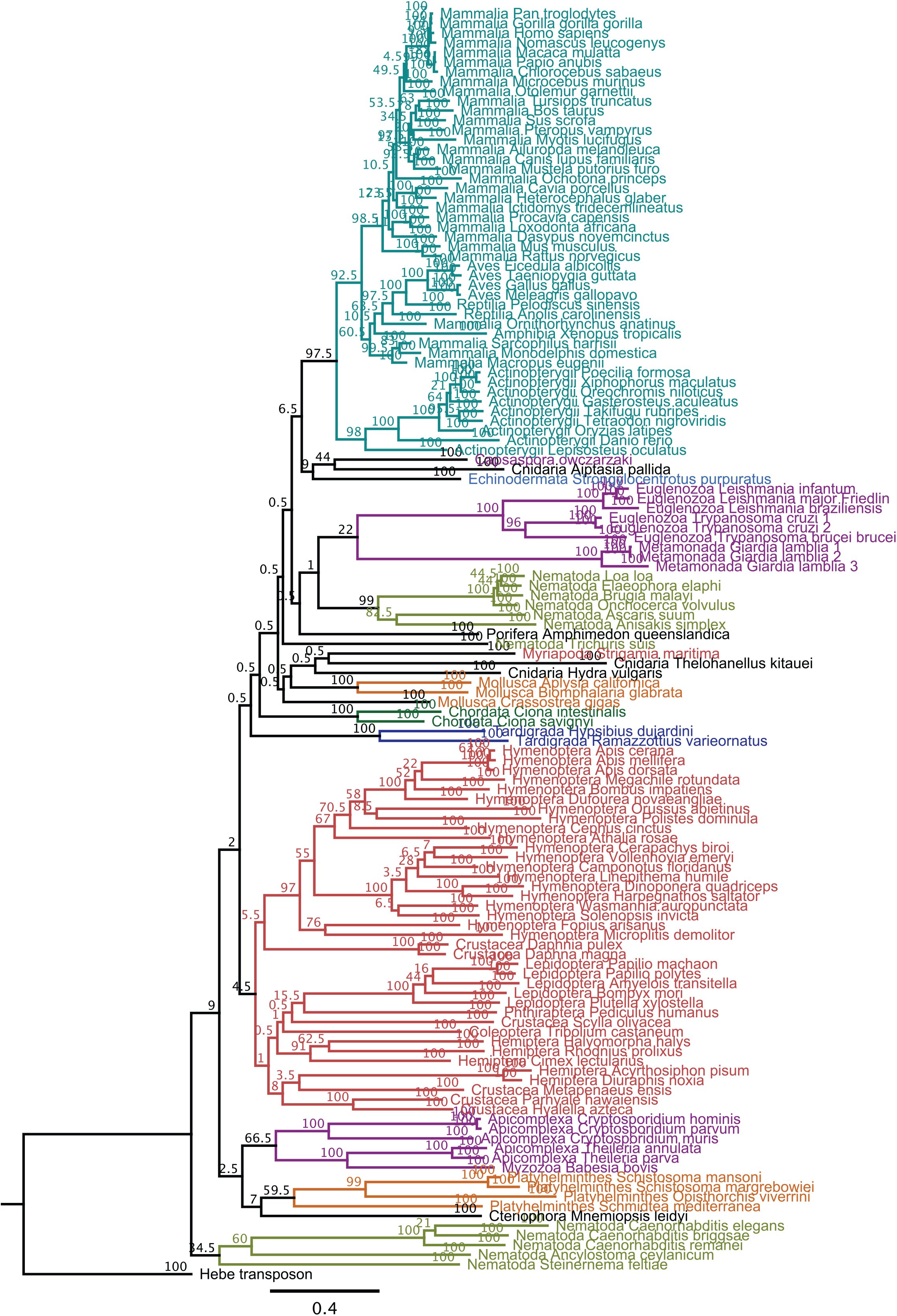
Phylogenetic tree of TERT sequences from representative metazoan phyla, early-branching metazoans and unicellular relatives of metazoans constructed using the maximum-likelihood method. Bootstrap support values from 1000 replicates are labelled at the nodes. Scale bar denotes substitutions per site.

**Supplementary Figure S2.**
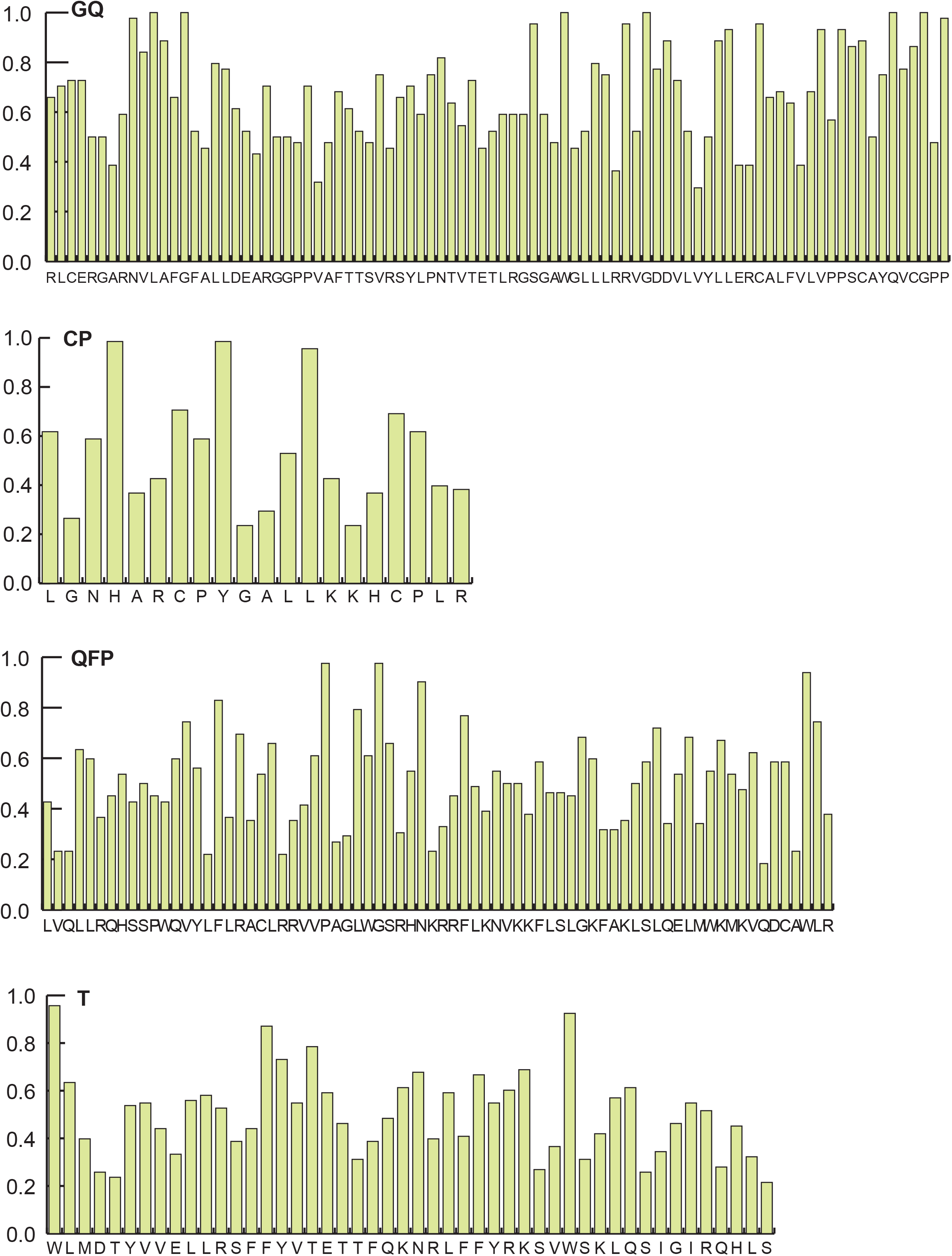
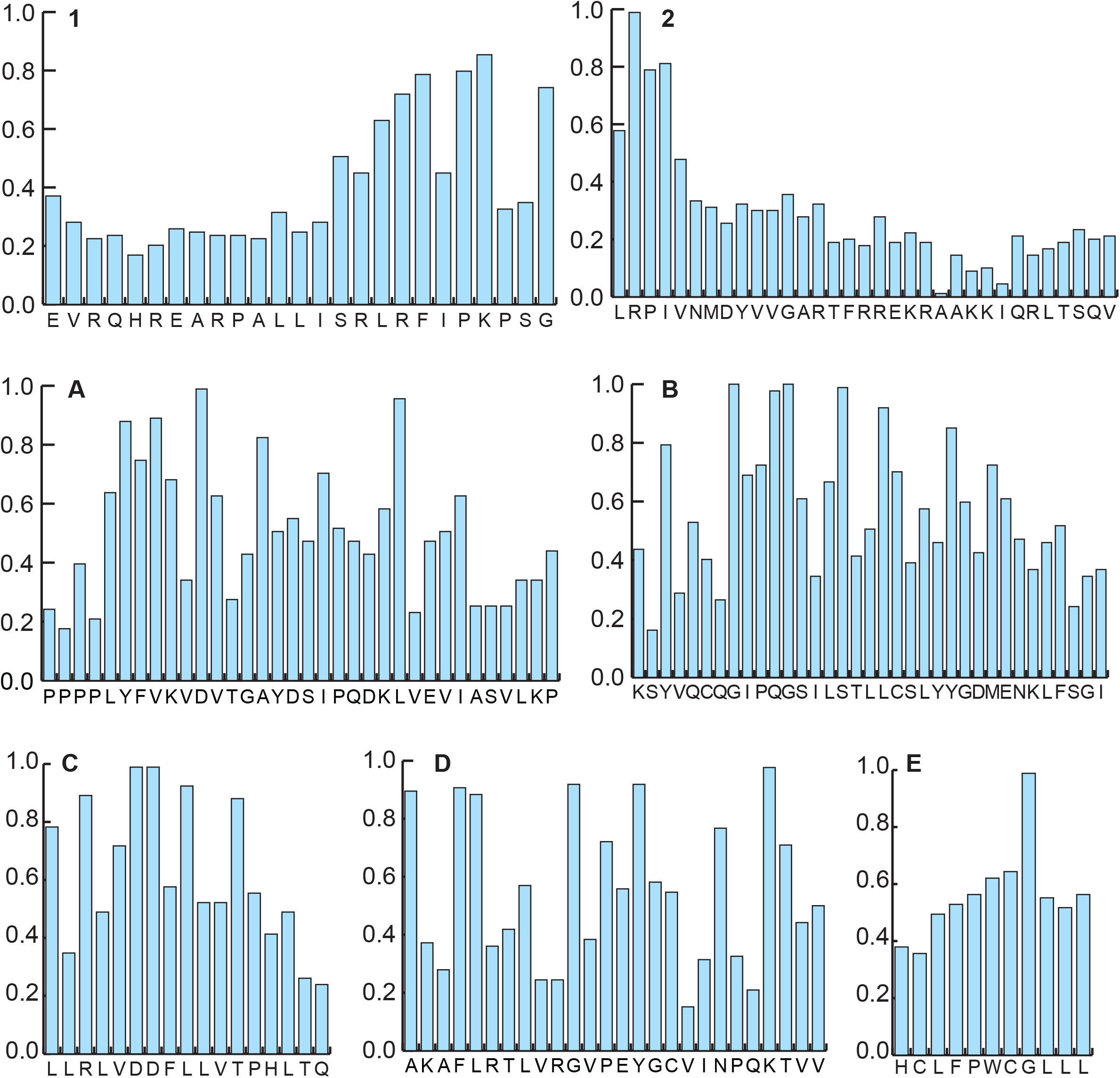
Graphs showing sequence conservation of TERT canonical motifs: TEN domain (GQ), TRBD domain (CP, QFP and T), and RT domain (1, 2, A, B’, C, D and E). The y-axes represent frequency of the most-abundant amino acid residue at any given position with a frequency of 1.0 indicating 100%. The x-axes represent the most-abundant amino acid residue at any given position across the length of the motif.

**Supplementary Figure S3.**
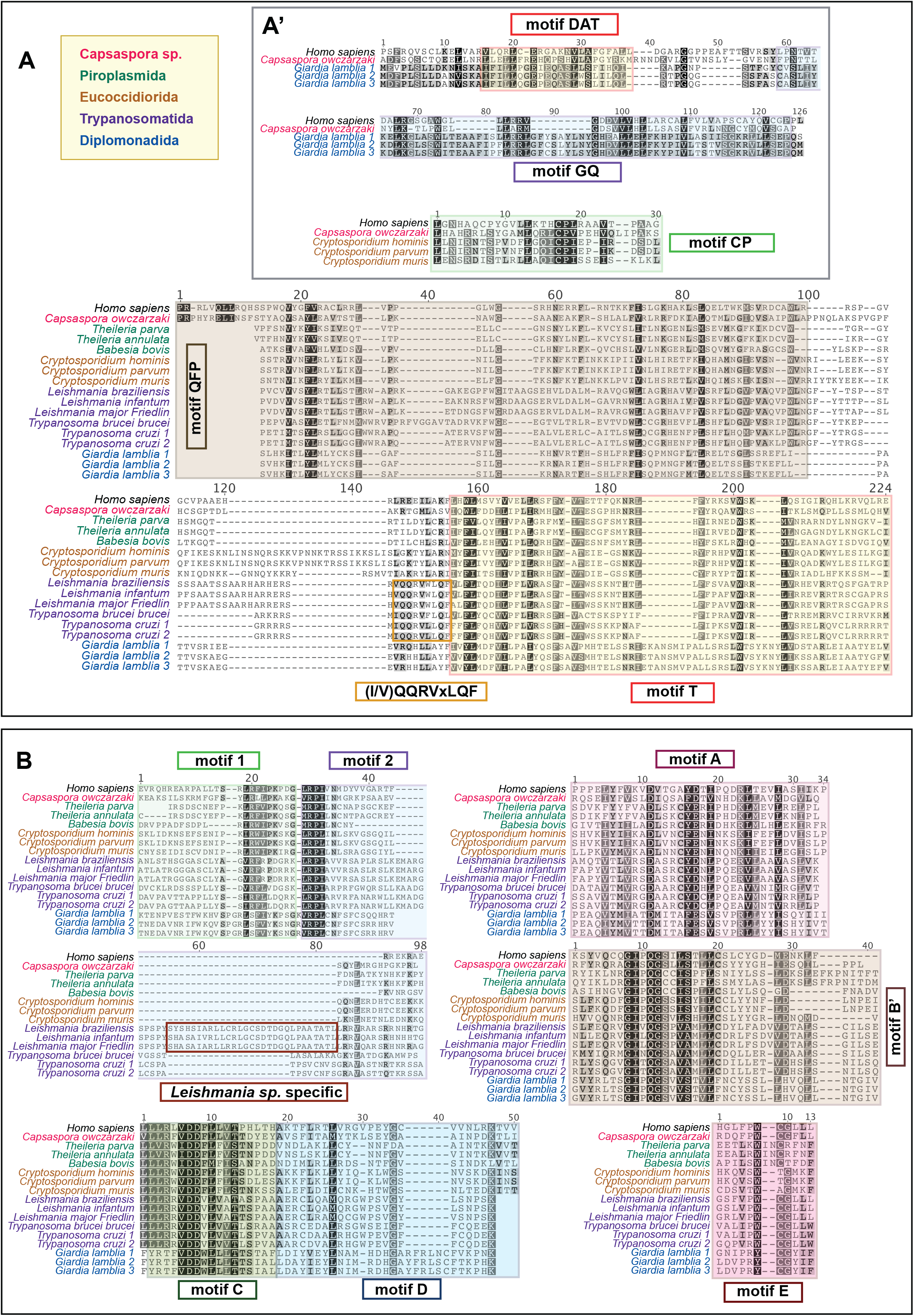
Multiple sequence alignments of the **(A)** TEN domain (GQ motif) and TRBD domain (CP, QFP, T) and **(B)** RT domain (1, 2, A, B’, C, D and E) from unicellular relatives of metazoans. Areas highlighted in coloured boxes delimit the canonical motifs. Boxes also denote species- or class-specific motifs with their respective descriptions written inside. Except for *Capsaspora*, the protozoan orders are represented by colour codes in the figure inset.

**Supplementary Figure S4.**
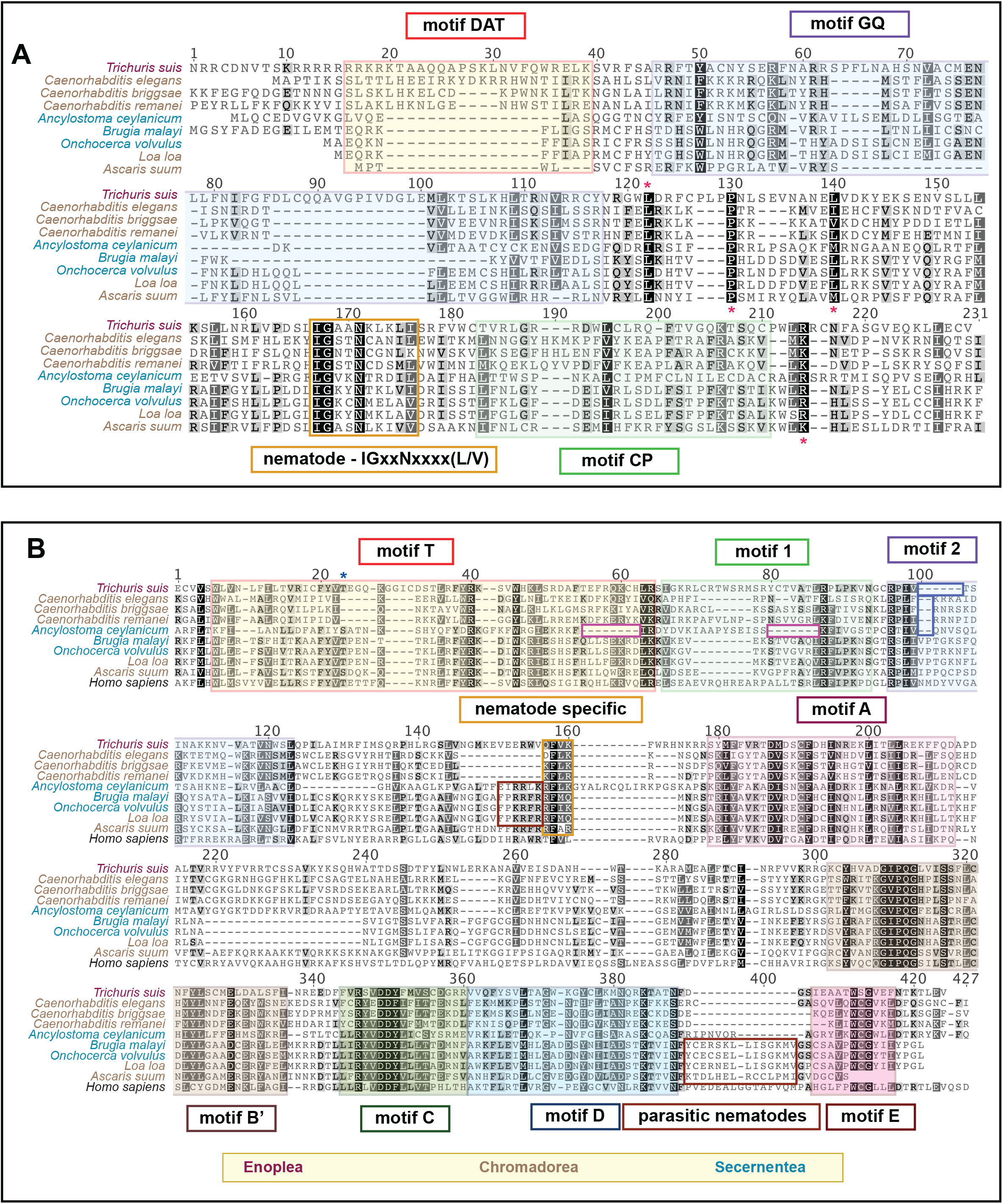
Multiple sequence alignments of the **(A)** GQ and CP motifs and **(B)** T motif and RT domain (1, 2, A, B’, C, D and E) from nematodes. Areas highlighted in coloured boxes delimit the canonical motifs. Boxes also denote Species- or Class-specific motifs with their respective descriptions written inside. Nematode classes are represented by colour codes in the figure inset.

**Supplementary Figure S5.**
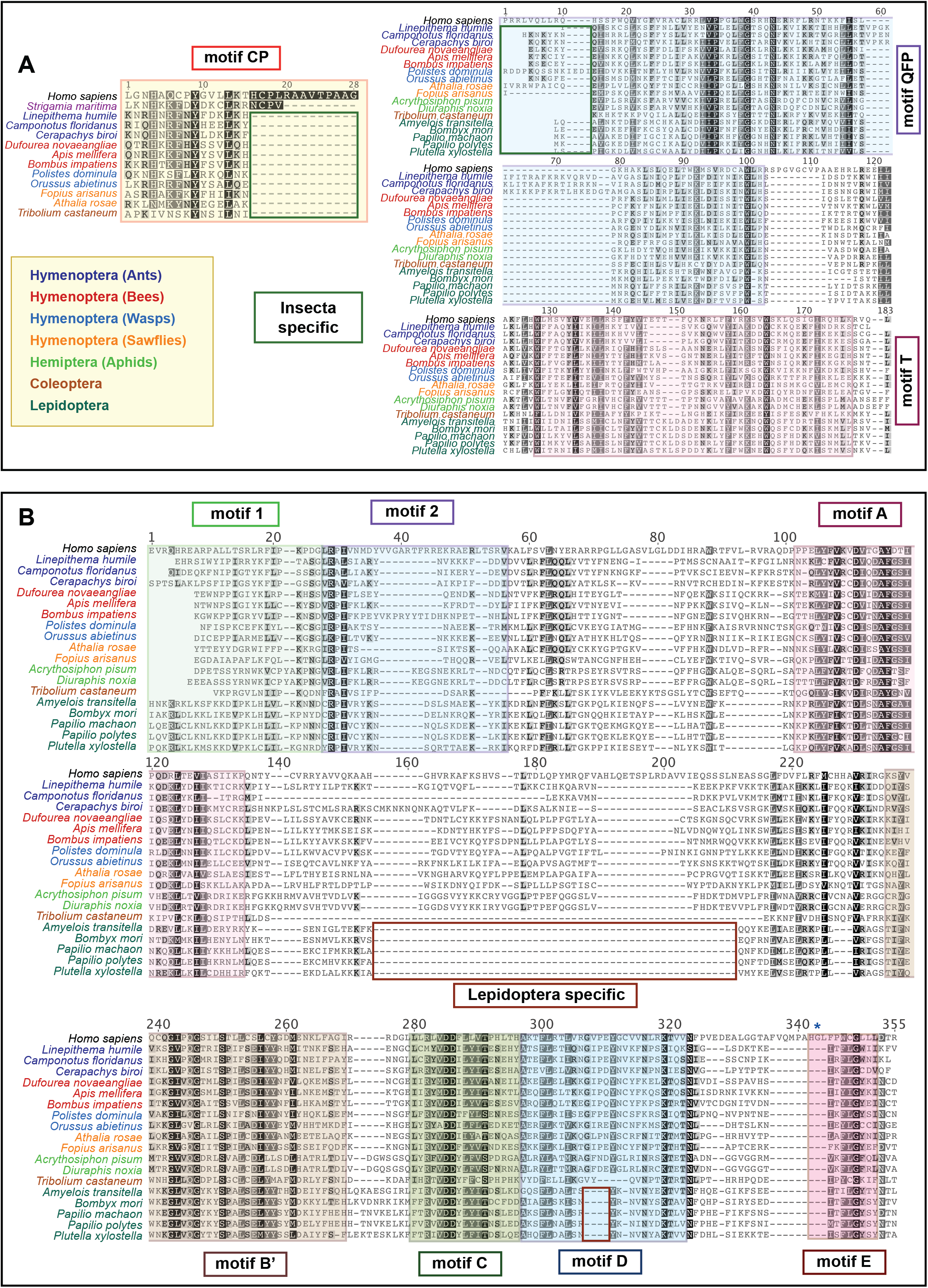
Multiple sequence alignments of the **(A)** TRBD domain (CP, QFP and T) and **(B)** and RT domain (1, 2, A, B’, C, D and E) from insects. Areas highlighted in coloured boxes delimit the canonical motifs. Boxes also denote Species- or order-specific motifs with their respective descriptions written inside. Insect orders are represented by colour codes in the figure inset.

**Supplementary Figure S6.**
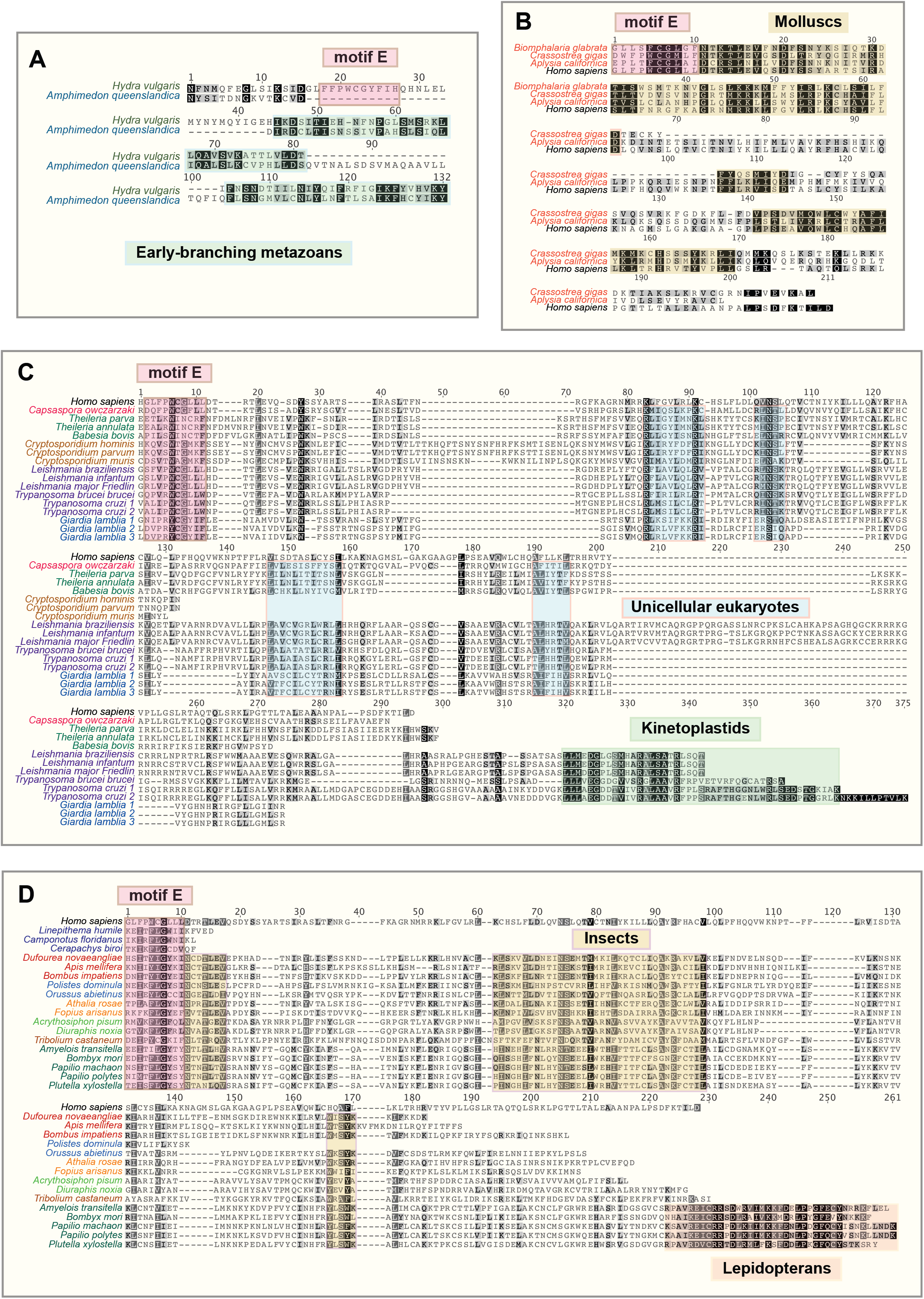
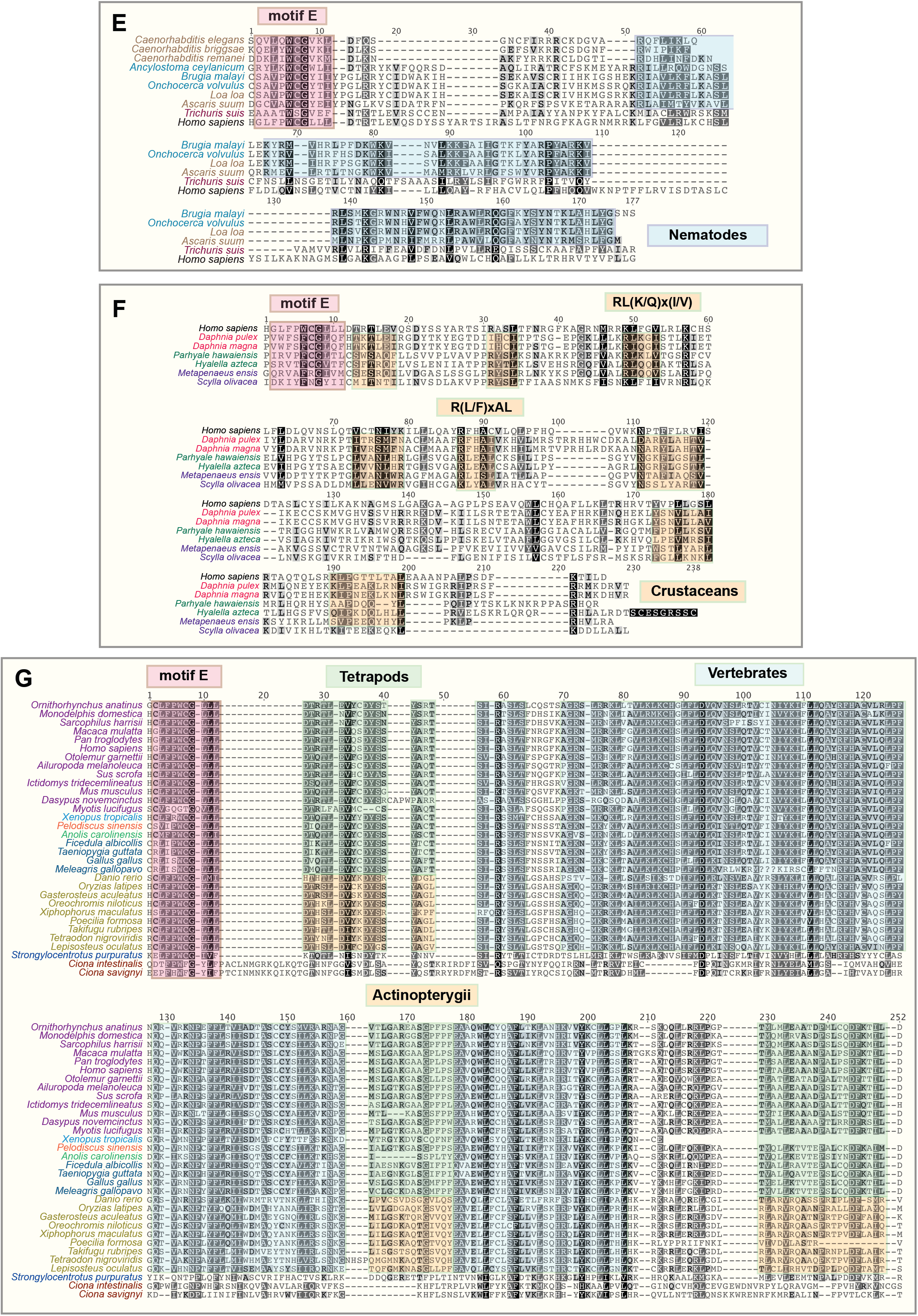
Multiple sequence alignments of the C-terminal extensions (CTEs) from (A) early-branching metazoans, (B) molluscs, (C) unicellular relatives of metazoans, (D) insects, (E) nematodes, (F) crustaceans and (G) vertebrates. The last TERT canonical motif E is highlighted in pink boxes to illustrate the start of CTE regions. Boxes also denote phylum-, order- or class-specific motifs with their respective descriptions written inside.

**Supplementary Figure S7.**
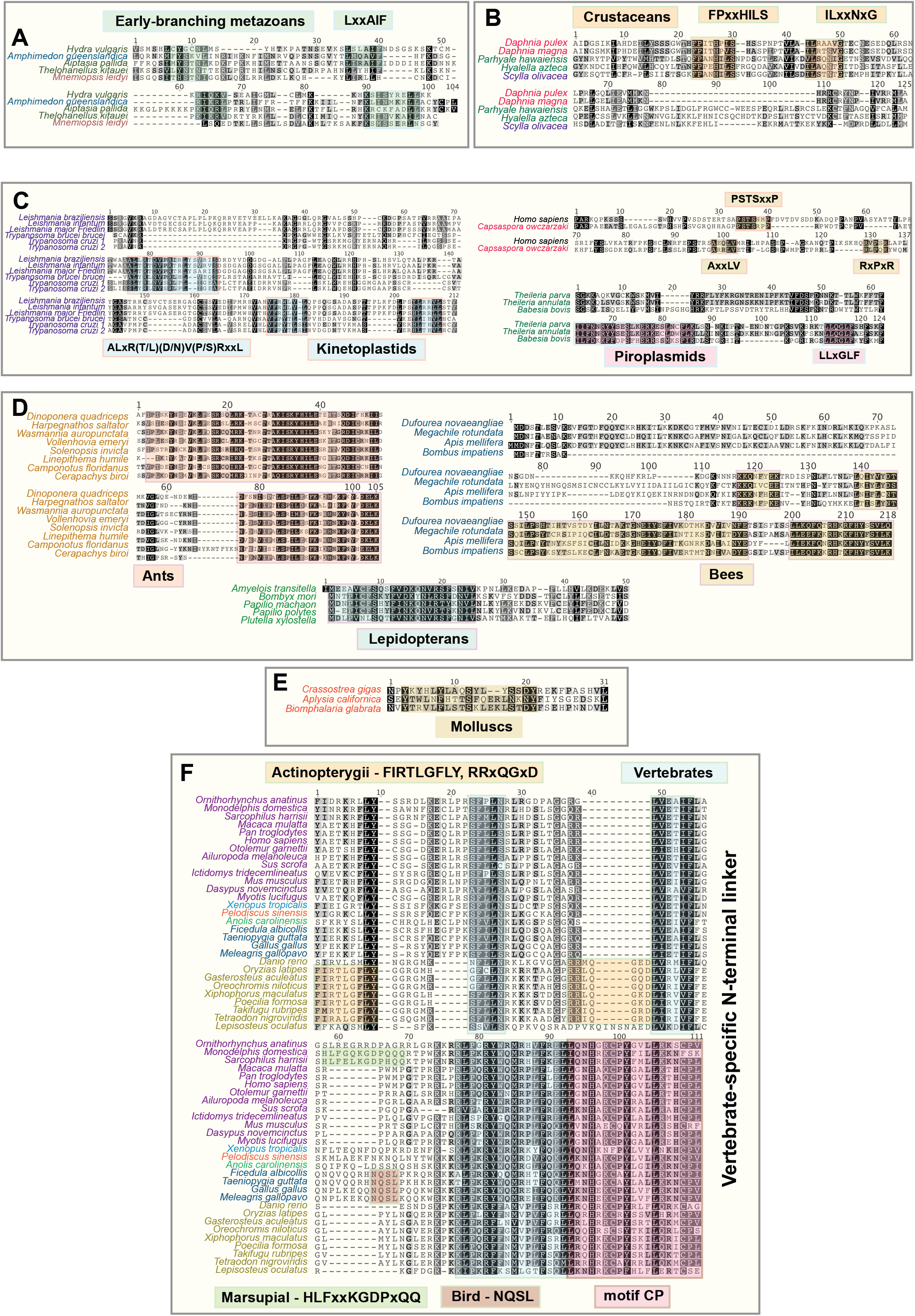
Multiple sequence alignments of the N-terminal linker regions from (A) early-branching metazoans, (B) crustaceans, (C) unicellular relatives of metazoans, (D) insects, (E) molluscs and (F) vertebrates. Boxes denote phylum-, order- or class-specific motifs with their respective descriptions written inside.

**Supplementary Figure S8.**
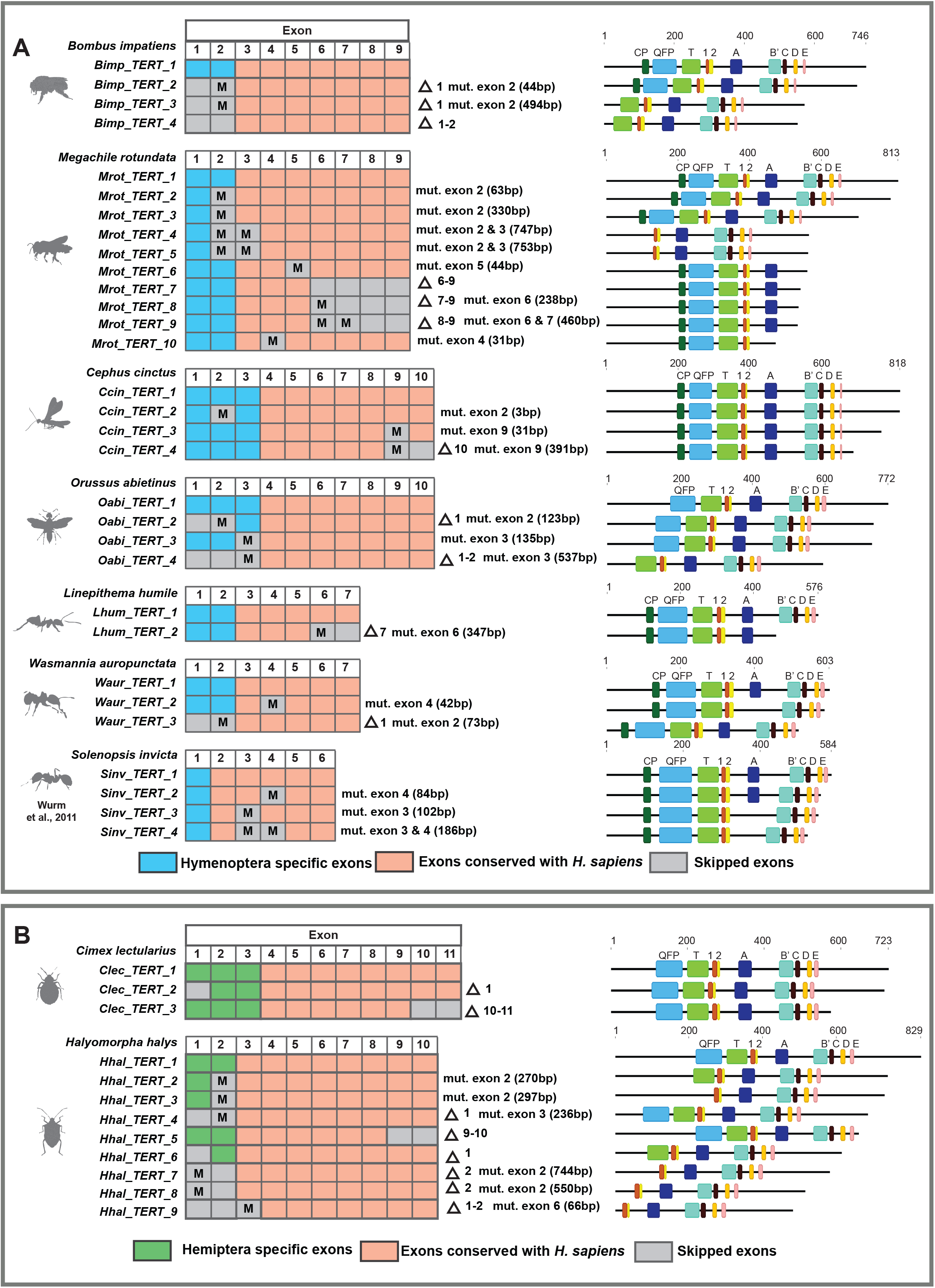
*TERT* alternative splicing in Hymenopteran and Hemipteran insects. *TERT* AS variants data were obtained from Ensembl Genomes or NCBI. These are generated by computational predictions based on RNA sequencing read evidence. Schematic diagram depicts gene structure and splicing of *TERT*. Orange boxes represent exons conserved with *hTERT*. Blue boxes represent Hymenoptera specific exons while green boxes represent Hemiptera specific exons. Gray boxes represent skipped or deleted exons resulted from alternative splicing events. ‘M’ denotes splice site mutations, deletions or intron retention. The left margin shows *TERT* gene and AS variant names for each species and the right margin shows descriptive names of *TERT* AS sequences. The numbers in parentheses represent the length of splice site mutations (deletion or intron retention) for the respective AS variants. Schematic diagrams on the far right illustrate the presence or absence of canonical motifs (CP, QFP, T, 1, 2, A, B’, C, D and E) on TERT AS protein variants drawn to scale.

**Supplementary Figure S9.**
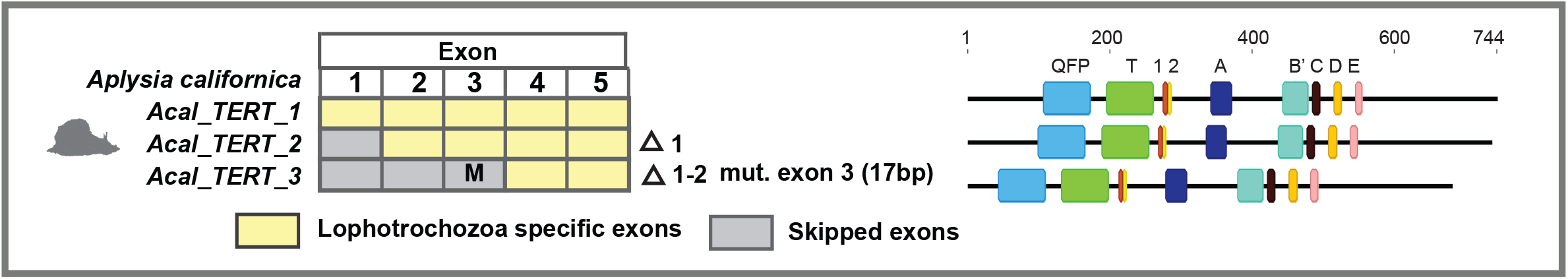
*TERT* alternative splicing in mollusc. Schematic diagram depicts gene structure and *TERT* AS variants. Yellow boxes represent *Aplysia californica* exons. Gray boxes represent skipped exons resulted from alternative splicing events. ‘M’ denotes splice site mutations, deletions or intron retention. The left margin shows *TERT* gene and AS variant names for each species and the right margin shows descriptive names of *TERT* AS sequences. The numbers in parentheses represent the length of splice site mutations (deletion or intron retention) for the respective AS variants. Schematic diagrams on the far right illustrate the presence or absence of canonical motifs (QFP, T, 1, 2, A, B’, C, D and E) on TERT AS protein variants drawn to scale.

**Supplementary Figure S10.**
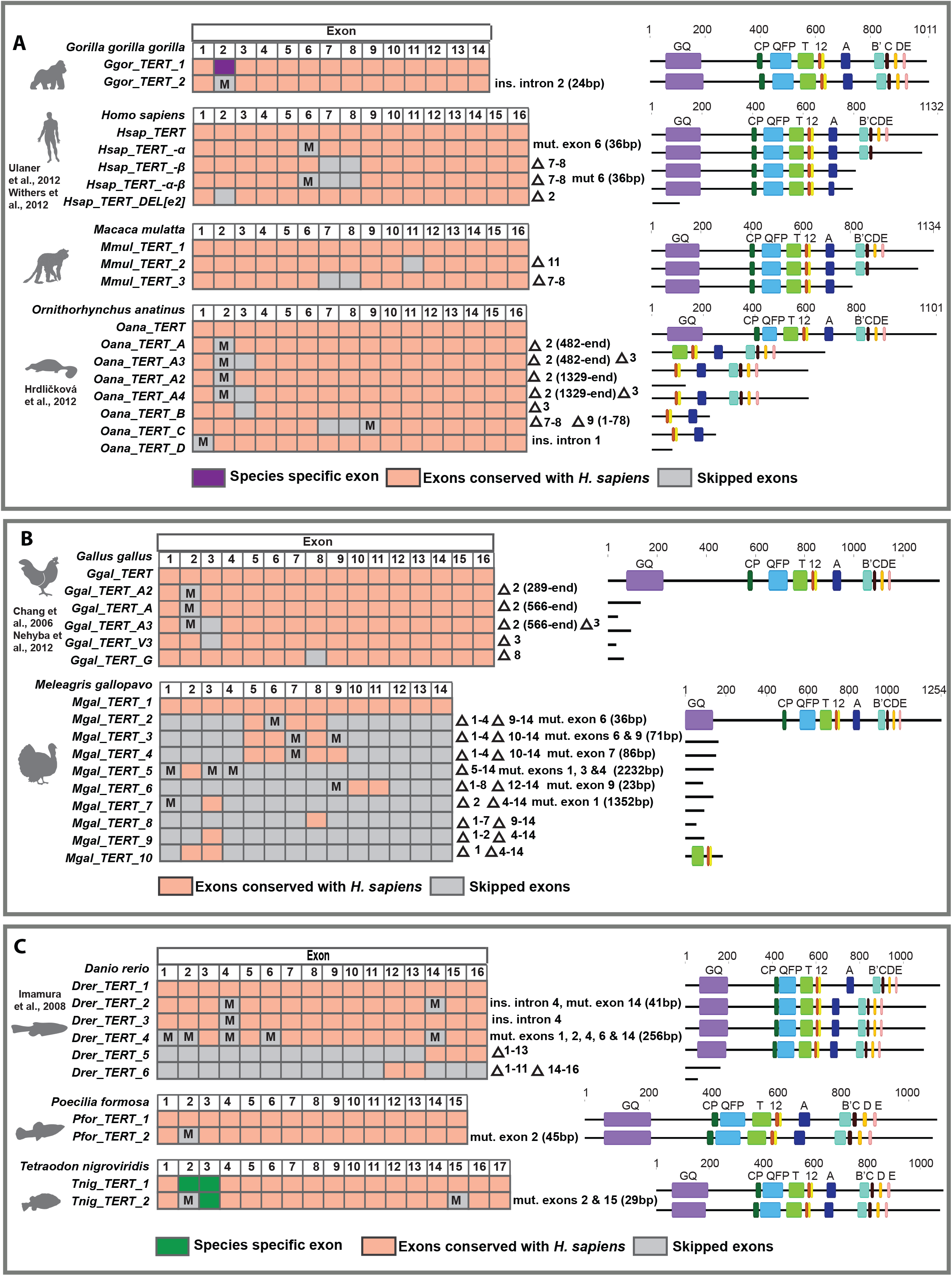
*TERT* alternative splicing in vertebrates. Only the main AS variants for *hTERT* and chicken *TERT* are shown in this diagram. Additional variants have been reported in Hrdličková et al. (2012). The presence of AS variants with skipped exon 2 is conserved across different vertebrate species. AS variants with spliced exons 7 to 8 are conserved in mammals except in *G. gorilla gorilla*. Mammalian *TERT* has 16 exons whereas *TERTs* in non-mammalian vertebrates have anywhere between 14 to 17 exons. *TERT* AS variants data was obtained from Ensembl Genomes or NCBI. These are generated by computational predictions based on RNA sequencing read evidence. Schematic diagram depicts gene structure and splicing of *TERT*. Orange boxes represent exons conserved with *hTERT*. Purple and green boxes represent species-specific exons. Grey boxes represent skipped or deleted exons resulted from alternative splicing events. ‘M’ denotes splice site mutations, deletions or intron retention. The left margin shows *TERT* gene and AS variant names for each species and the right margin shows descriptive names of *TERT* AS sequences. The numbers in parentheses represent the length of splice site mutations (deletion or intron retention) for the respective AS variants except for platypus (*O. anatinus*) and chicken (*G. gallus*). For these two species, the numbers in parentheses represent the position of deletion in the exon indicated. Schematic diagrams on the far right illustrate the presence or absence of canonical motifs (GQ, CP, QFP, T, 1, 2, A, B’, C, D and E) on TERT AS protein variants drawn to scale.

**Supplementary figure S11.**
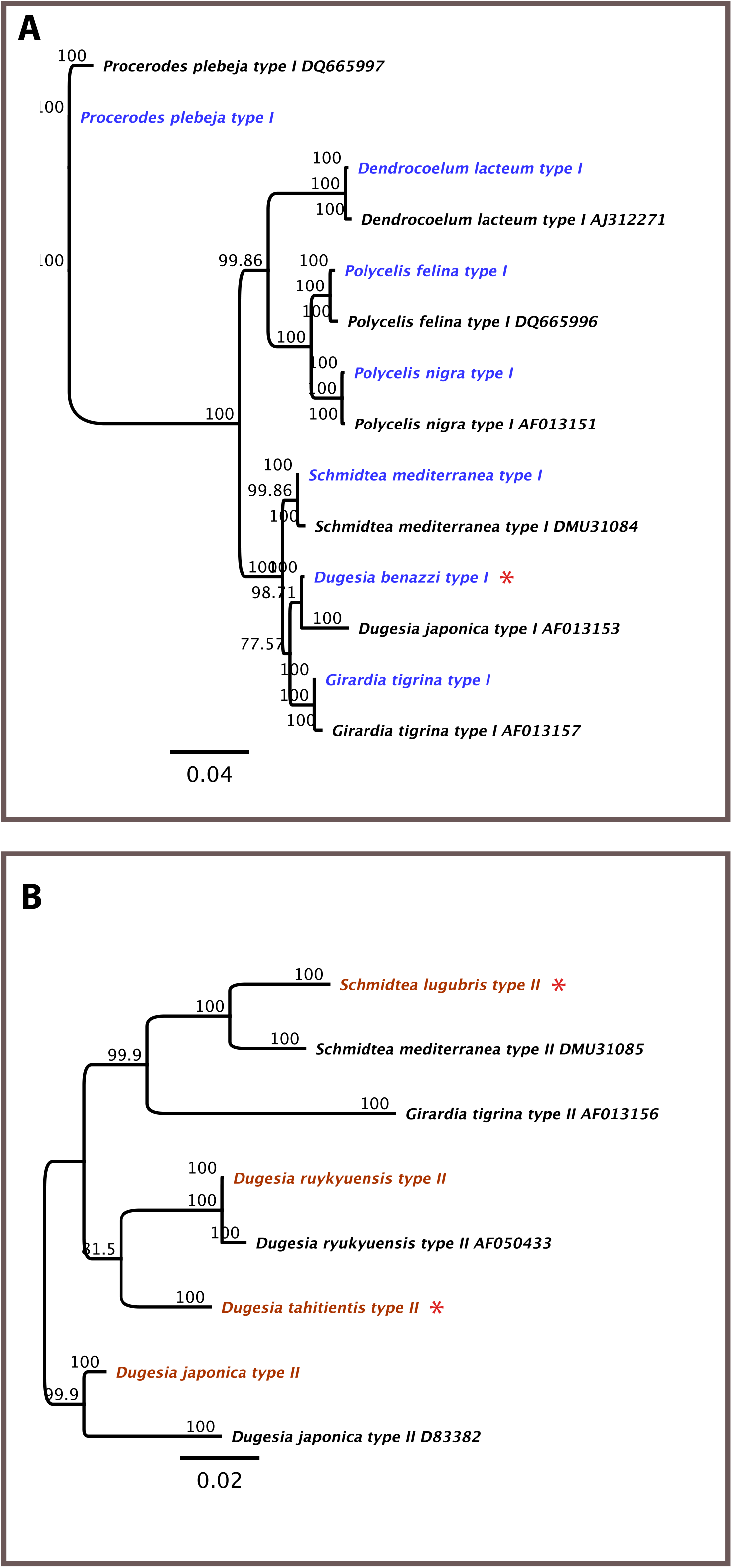
Phylogenetic tree illustrating the positions of planarian type I and type II sequences in the 18S rDNA phylogeny. Sequences of 18S rDNA were cloned from 11 flatworm species in the lab for comparison against published sequences in GenBank. **(A)** Type I 18S rDNA sequences with species marked in blue as cloned in the lab. **(B)** Type II 18S rDNA sequences with species marked in brown as cloned in the lab. Asterisks indicate novel 18S rDNA sequences obtained from this study.

**Supplementary figure S12.**
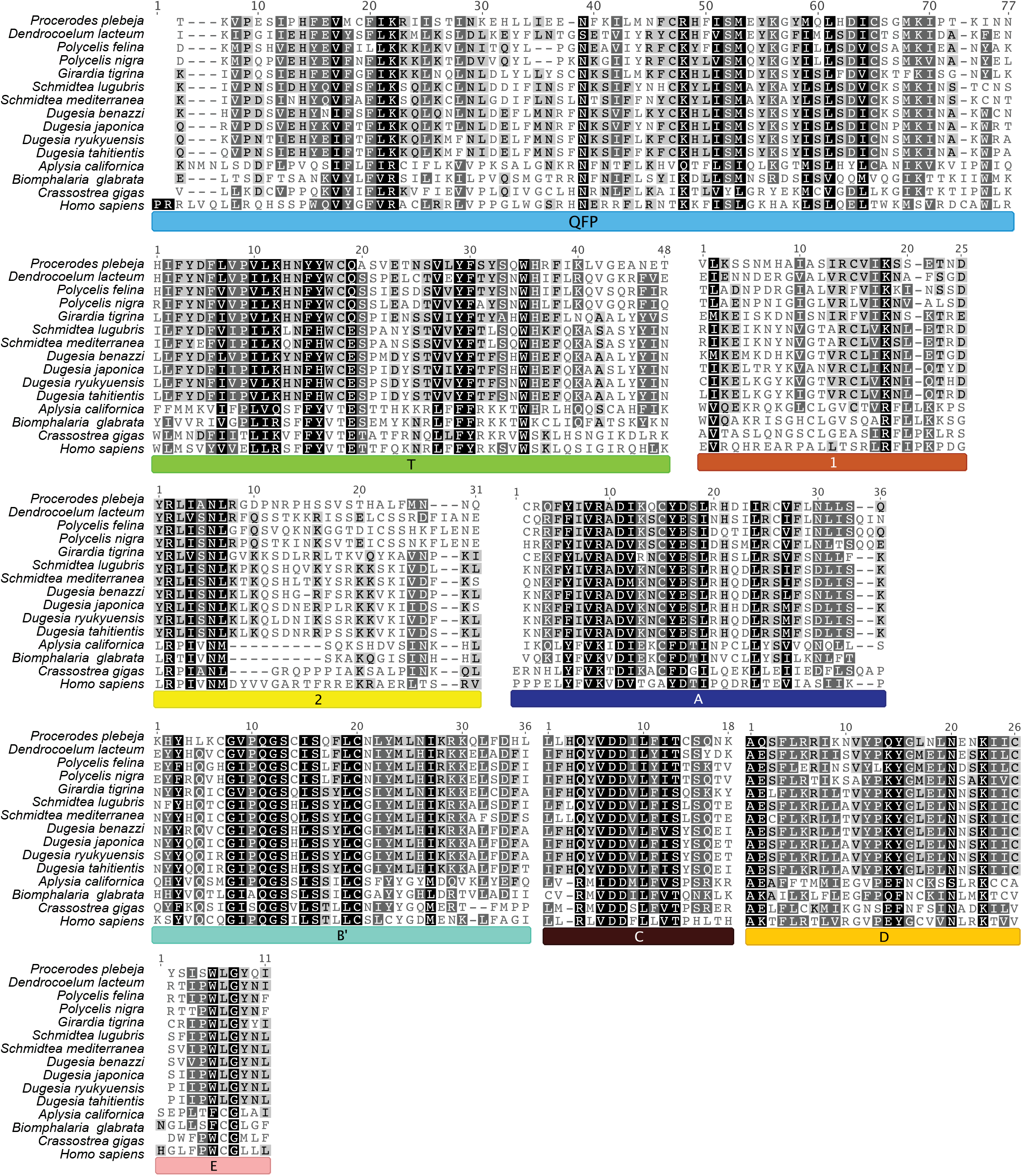
Alignment of canonical motifs of TERT from 11 planarian species, three molluscs and hTERT. These include TRBD domain (QFP and T motifs) and canonical reverse transcriptase motifs (1, 2, A, B’, C, D and E). Clearly defined GQ and CP motifs were not identified from planarian TERTs.

**Supplementary figure S13.**
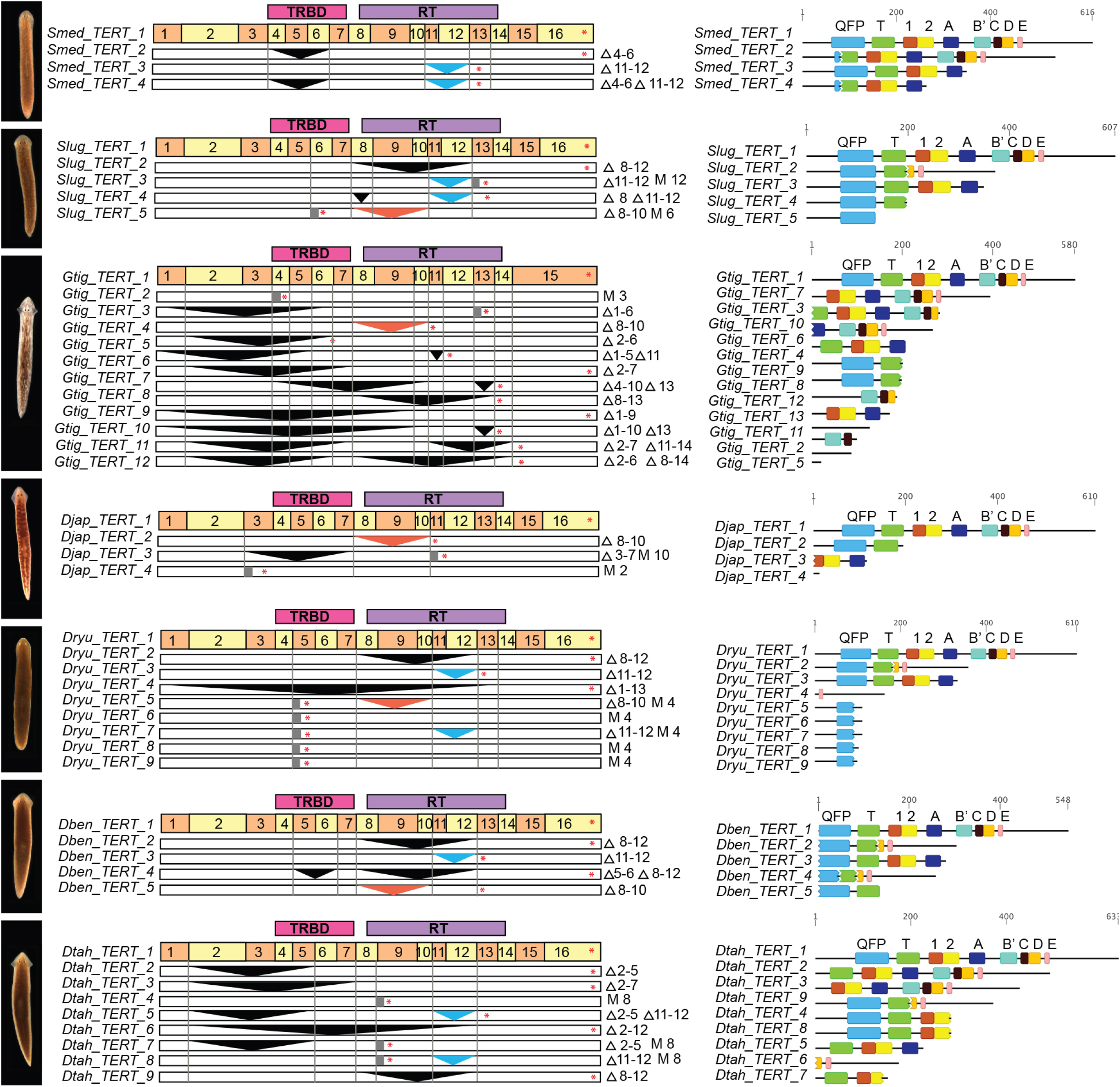
*TERT* in Dugesiidae is alternatively spliced. Comparison of alternatively spliced (AS) variants in seven Dugesiidae species from the genera *Schmidtea, Girardia and Dugesia*. Structure of AS variants in *S. mediterranea, S. lugubris, G. tigrina, D. japonica, D. ryukyuensis, D. benazzi and D. tahitientis*. Full-length or wild-type *TERT* structure is shown at the top of each set of AS variants. Functional TERT domains (TRBD and RT) and positions of (putative) exons are indicated. Deletions (skipped exons) are denoted by triangles and insertions (retained introns or splice site mutations) are denoted by grey rectangles. 'M' abbreviations represent splice site mutations. Red triangles represent conserved alternatively spliced exons in all Dugesiidae species except *S. mediterranea* and *D. tahitientis*. Blue triangles represent conserved alternatively spliced exons in all Dugesiidae species except *G. tigrina* and *D. japonica*. Asterisks indicate stop codon positions caused by frame shift mutation or retained introns. The left margin shows *TERT* gene and AS variant names for each species and the right margin shows descriptive names of *TERT* AS sequences. Schematic diagrams on the far right illustrate the presence or absence of canonical motifs (QFP, T, 1, 2, A, B’, C, D and E) on TERT AS protein variants drawn to scale.

**Supplementary figure S14.**
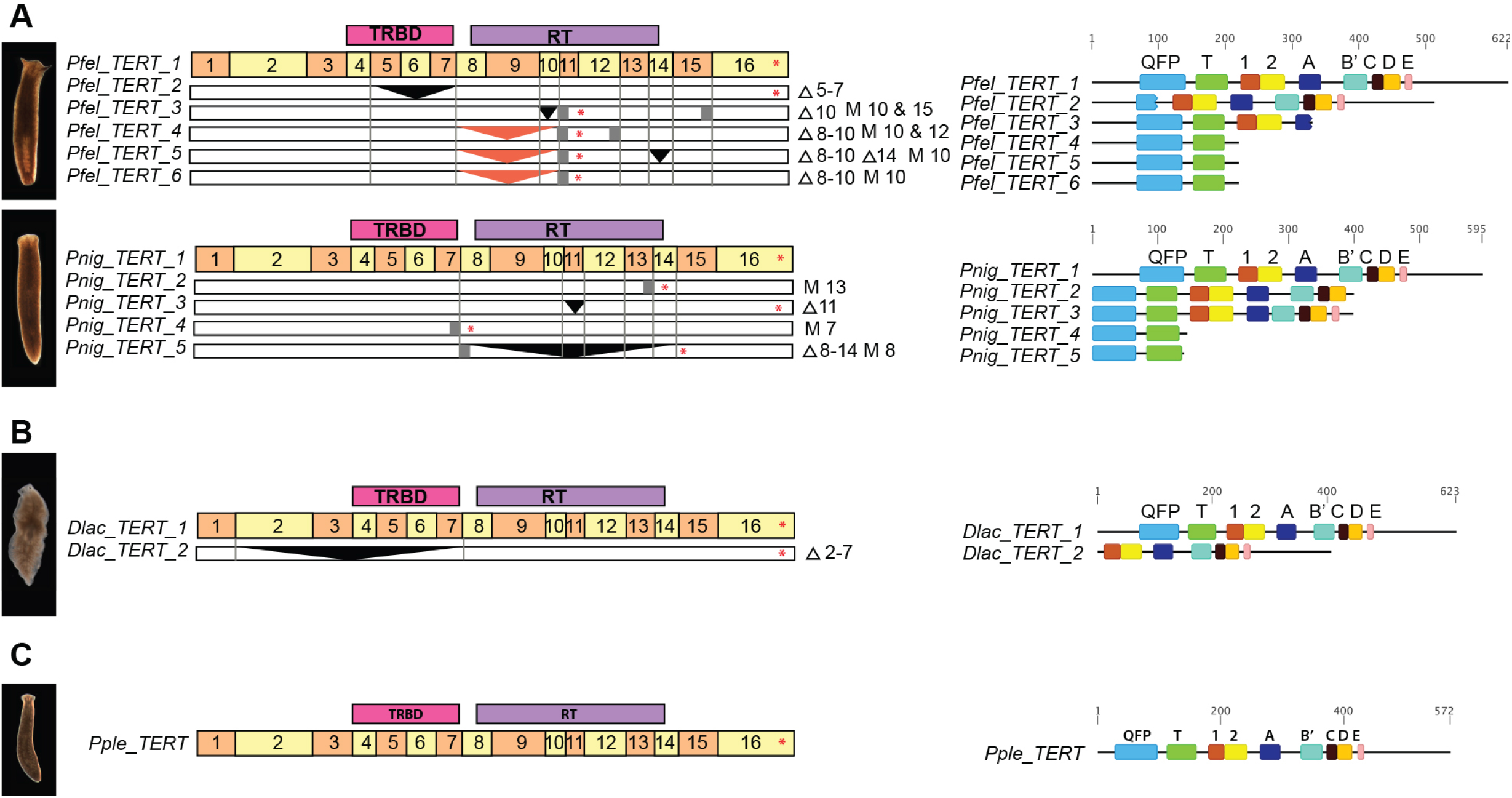
*TERT* in Planariidae and Dendrocoelidae but not Procerodidae is alternatively spliced. Comparison of AS variants in two Planariidae species from the genus *Polycelis*, one Dendrocoelidae species and one Procerodidae species. Structure of AS variants in **(A)** Planariidae *P. felina* and *P. nigra*, **(B)** Dendrocoelidae *D. lacteum* and **(C)** Procerodidae *P. plebeja*. Genomic sequence data is not available for these flatworm species. Therefore, putative *TERT* exon-intron boundaries were annotated based on *Smed_TERT* sequence and confirmed by AS variant analyses of each species. Full-length or wild-type *TERT* structure is shown at the top of each set of AS variants. Functional TERT domains (TRBD and RT) and positions of (putative) exons are indicated. Deletions (skipped exons) are denoted by triangles and insertions (retained introns or splice site mutations) are denoted by gray rectangles. Red triangles represent alternatively spliced exons conserved with Dugesiidae. Asterisks indicate stop codon positions caused by frame shift mutation or retained introns. The left margin shows *TERT* gene and AS variant names for each species and the right margin shows descriptive names of *TERT* AS sequences. Schematic diagrams on the far right illustrate the presence or absence of canonical motifs (QFP, T, 1, 2, A, B’, C, D and E) on TERT AS protein variants drawn to scale.

**Supplementary figure S15.**
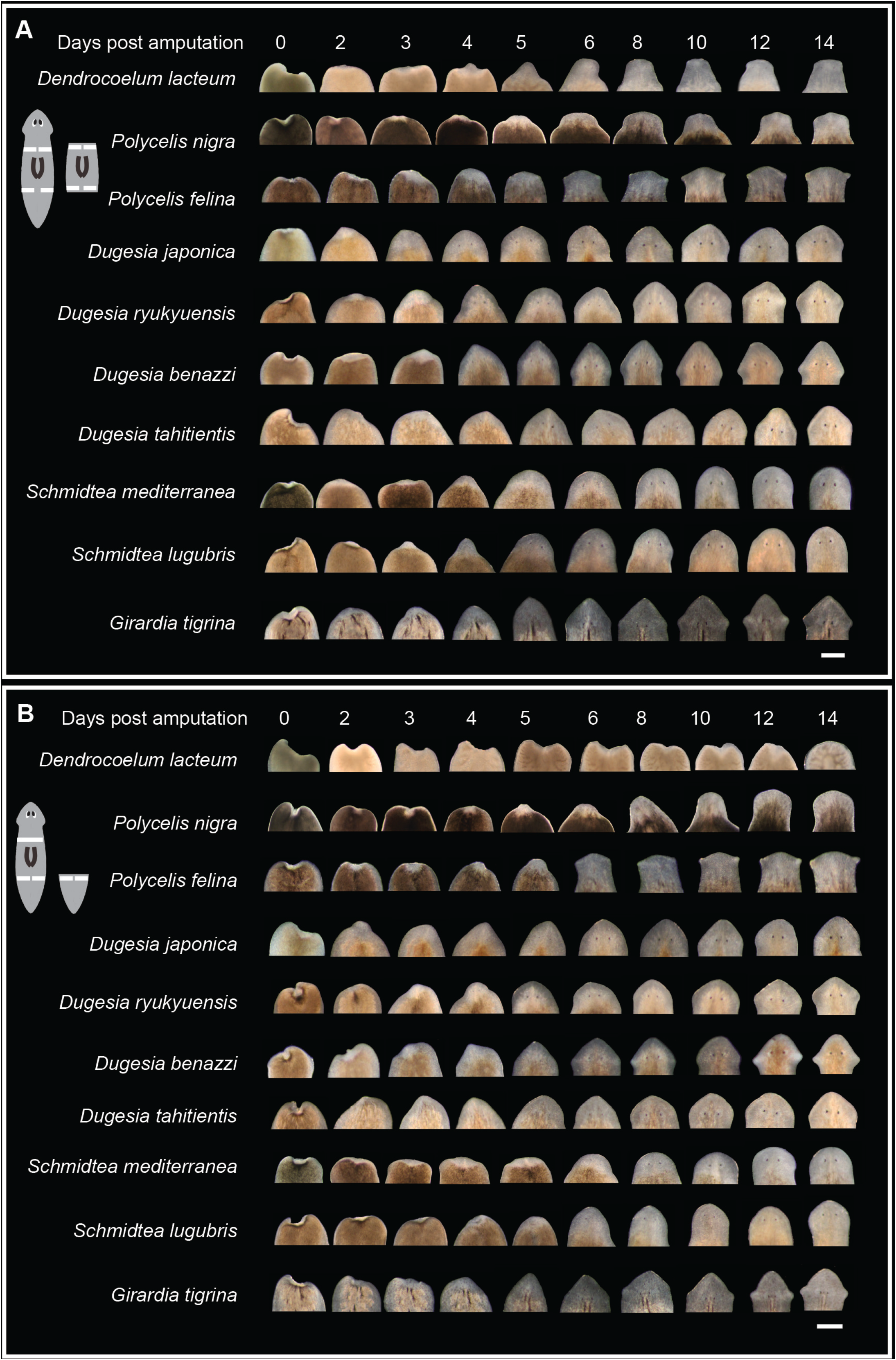
Regenerative capacity in planarians. Transverse sections assaying regenerative abilities along anterior-posterior axis of planarians. Ten worms were used for each species and were amputated into three fragments each (head, trunk and tail). Representative images depict time-course observation of the same animal undergoing cephalic regeneration in **(A)** trunk and **(B)** tail fragments. Anterior blastema (unpigmented region) was imaged. Appearances of pigmented eyes were noted and plotted as frequency graphs. Scale bars indicate 0.5mm.

**Supplementary figure S16.**
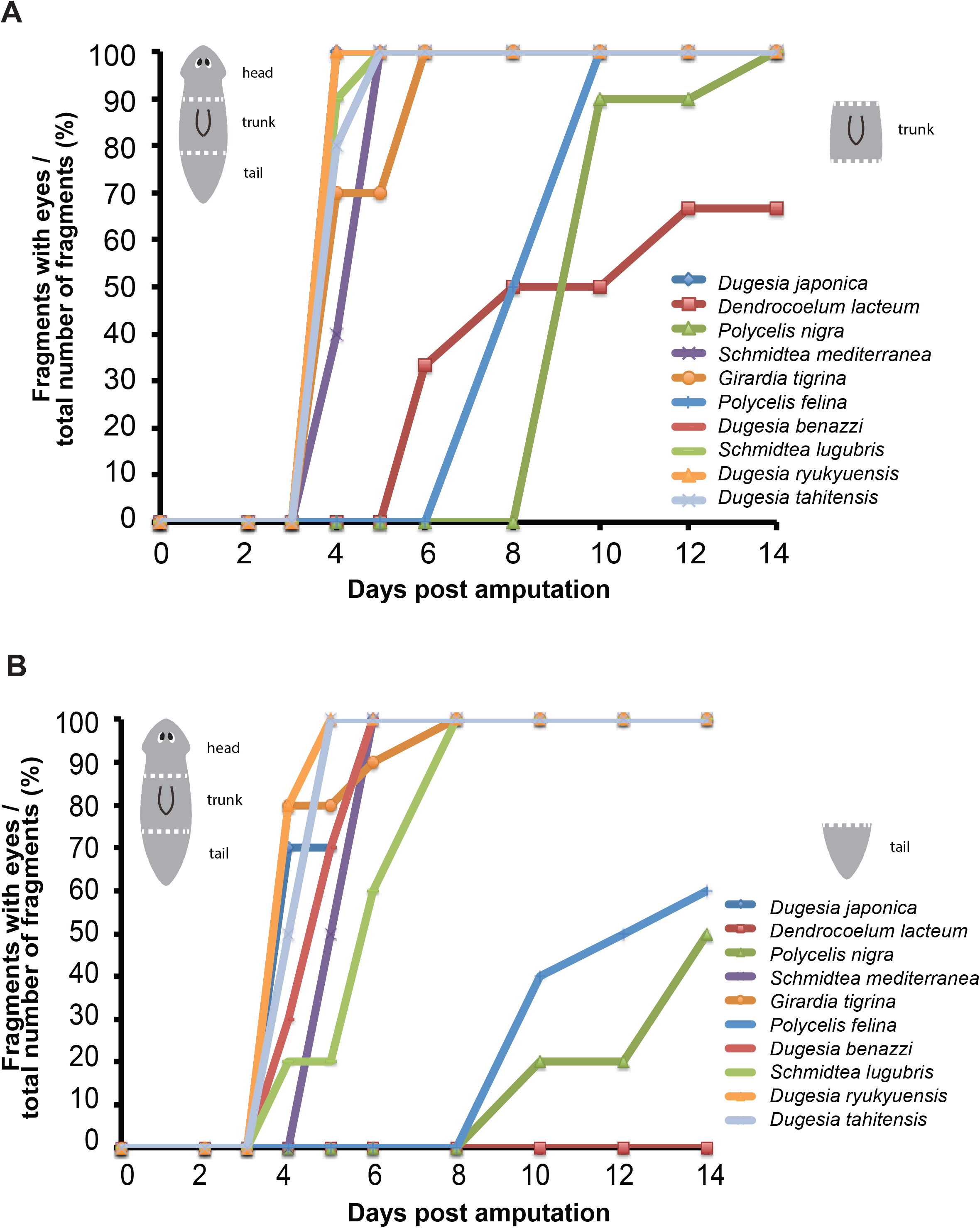
Head regeneration frequency graph for **(A)** trunk fragments and **(B)** tail fragments of amputated flatworms (N=10 for each species). Cartoon depicts the amputation planes used. The timing of eye appearance in cohorts of 10 fragments was noted. *P. plebeja* failed to regenerate at all and is not shown.

## References

Abad JP, De Pablos B, Osoegawa K, De Jong PJ, Martín-Gallardo A, Villasante A. 2004. *TAHRE*, a novel telomeric retrotransposon from *Drosophila melanogaster*, reveals the origin of Drosophila telomeres. Mol Biol Evol. 21:1620–1624.

Adam RD, Dahlstrom EW, Martens CA, Bruno DP, Barbian KD, Ricklefs SM, Hernandez MM, Narla NP, Patel RB, Porcella SF, et al. 2013. Genome sequencing of *Giardia lamblia* genotypes A2 and B Isolates (DH and GS) and comparative analysis with the genomes of genotypes A1 and E (WB and pig). Genome Biol and Evol. 5:2498–2511.

Aken BL, Ayling S, Barrell D, Clarke L, Curwen V, Fairley S, Fernandez Banet J, Billis K, García Girón C, Hourlier T, et al. 2016. The Ensembl gene annotation system. Database 2016:baw093.

Armbruster BN, Banik SS, Guo C, Smith AC, Counter CM. 2001. N-terminal domains of the human telomerase catalytic subunit required for enzyme activity in vivo. Mol Cell Biol. 21:7775–7786.

Autexier C, Greider CW. 1995. Boundary elements of the Tetrahymena telomerase RNA template and alignment domains. Genes Dev. 9:2227–2239.

Autexier C, Lue NF. 2006. The structure and function of telomerase reverse transcriptase. Annu Rev Biochem. 75:493–517.

Álvarez-Presas M, Baguñà J, Riutort M. 2008. Molecular phylogeny of land and freshwater planarians (Tricladida, Platyhelminthes): From freshwater to land and back. Molecular Phylogenet Evol. 47:555–568.

Baumgarten S, Simakov O, Esherick LY, Liew YJ, Lehnert EM, Michell CT, Li Y, Hambleton EA, Guse A, Oates ME, et al. 2015. The genome of *Aiptasia*, a sea anemone model for coral symbiosis. Proc Natl Acad Sci U.S.A. 112:11893–11898.

Beattie TL, Zhou W, Robinson MO, Harrington L. 2001. Functional multimerization of the human telomerase reverse transcriptase. Mol Cell Biol. 21:6151–6160.

Biessmann H, Carter SB, Mason JM. 1990. Chromosome ends in Drosophila without telomeric DNA sequences. Proc Natl Acad Sci U.S.A. 87:1758–1761.

Biessmann H, Champion LE, O'Hair M, Ikenaga K, Kasravi B, Mason JM. 1992. Frequent transpositions of Drosophila melanogaster *HeT-A* transposable elements to receding chromosome ends. EMBO J. 11:4459–4469.

Bischoff C, Petersen HC, Graakjaer J, Andersen-Ranberg K, Vaupel JW, Bohr VA, K lvraa S, Christensen K. 2006. No Association Between Telomere Length and Survival Among the Elderly and Oldest Old. Epidemiology 17:190–194.

Blackburn EH, Gall JG. 1978. A tandemly repeated sequence at the termini of the extrachromosomal ribosomal RNA genes in *Tetrahymena*. J Mol Biol. 120:33–53.

Blackburn EH, Greider CW, Szostak JW. 2006. Telomeres and telomerase: the path from maize, *Tetrahymena* and yeast to human cancer and aging. Nat Med. 12:1133–1138.

Blasco MA, Lee H-W, Hande MP, Samper E, Lansdorp PM, DePinho RA, Greider CW. 1997. Telomere Shortening and Tumor Formation by Mouse Cells Lacking Telomerase RNA. Cell 91:1–10.

Blasco MA. 2005. Telomeres and human disease: ageing, cancer and beyond. Nat Rev Genet. 6:611–622.

Bley CJ, Qi X, Rand DP, Borges CR, Nelson RW, Chen JJL. 2011. RNA-protein binding interface in the telomerase ribonucleoprotein. Proc Natl Acad Sci U.S.A. 108:20333–20338.

Brandl H, Moon H, Vila-Farré M, Liu S-Y, Henry I, Rink JC. 2016. PlanMine--a mineable resource of planarian biology and biodiversity. Nucleic Acids Res. 44:D764–D773.

Bryan TM, Englezou A, Gupta J, Bacchetti S, Reddel RR. 1995. Telomere elongation in immortal human cells. EMBO J. 14:4240–4248.

Bryan TM, Sperger JM, Chapman KB, Cech TR. 1998. Telomerase reverse transcriptase genes identified in *Tetrahymena thermophila* and *Oxytricha trifallax*. Proc Natl Acad Sci U.S.A. 95:8479–8484.

Cai Y, Ai Y, Zhao Q, Li J, Yang G, Gong P, Wang Q, Hou H, Zhang G, Li L, et al. 2011. Cloning and characterization of telomerase reverse transcriptase gene in *Trichinella spiralis*. Parasitol Res. 110:411–417.

Carranza S, Giribet G, Ribera C, Baguñà, Riutort M. 1996. Evidence that two types of 18S rDNA coexist in the genome of *Dugesia* (*Schmidtea*) *mediterranea* (Platyhelminthes, Turbellaria, Tricladida). Mol Biol Evol. 13:824–832.

Casacuberta E, Pardue ML. 2002. Coevolution of the telomeric retrotransposons across *Drosophila* species. Genetics 161:1113–1124.

Casacuberta E, Pardue ML. 2003. Transposon telomeres are widely distributed in the *Drosophila* genus: *TART* elements in the virilis group. Proc Natl Acad Sci U.S.A. 100:3363–3368.

Chang H, Delany ME. 2006. Complicated RNA splicing of chicken telomerase reverse transcriptase revealed by profiling cells both positive and negative for telomerase activity. Gene 379:33–39.

Chapman JA, Kirkness EF, Simakov O, Hampson SE, Mitros T, Weinmaier T, Rattei T, Balasubramanian PG, Borman J, Busam D, et al. 2010. The dynamic genome of *Hydra*. Nature 464:592–596.

Charnov EL, Berrigan D. 1991. Evolution of life history parameters in animals with indeterminate growth, particularly fish. Evol Ecol 5:63–68.

Chen J-L, Greider CW. 2003. Template boundary definition in mammalian telomerase. Genes Dev. 17:2747–2752.

Choi J, Southworth LK, Sarin KY, Venteicher AS, Ma W, Chang W, Cheung P, Jun S, Artandi MK, Shah N, et al. 2008. TERT Promotes Epithelial Proliferation through Transcriptional Control of a Myc- and Wnt-Related Developmental Program. PLoS Genet 4:e10–e15.

Colbourne JK, Pfrender ME, Gilbert D, Thomas WK, Tucker A, Oakley TH, Tokishita S, Aerts A, Arnold GJ, Basu MK, et al. 2011. The Ecoresponsive Genome of *Daphnia pulex*. Science 331:555–561.

Collins K, Gandhi L. 1998. The reverse transcriptase component of the *Tetrahymena* telomerase ribonucleoprotein complex. Proc Natl Acad Sci U.S.A. 95:8485–8490.

Collins K. 1999. Ciliate telomerase biochemistry. Annu Rev Biochem. 68:187–218.

Counter CM, Avilion AA, Lefeuvre CE, Stewart NG, Greider CW, Harley CB, Bacchetti S. 1992. Telomere shortening associated with chromosome instability is arrested in immortal cells which express telomerase activity. EMBO J. 11:1921–1929.

De Jesus BB, Vera E, Schneeberger K, Tejera AM, Ayuso E, Bosch F, Blasco MA. 2012. Telomerase gene therapy in adult and old mice delays aging and increases longevity without increasing cancer. EMBO Mol Med 4:691–704.

De Lange T. 2005. Telomere-related genome instability in cancer. Cold Spring Harbor Symposia on Quantitative Biology 70:197–204.

Dreesen O, Li B, Cross GAM. 2005. Telomere structure and shortening in telomerase-deficient Trypanosoma brucei. Nucleic Acids Res. 33:4536–4543.

Ebert TA, Russell MP, Gamba G, Bodnar A. 2008. Growth, survival, and longevity estimates for the rock-boring sea urchin *Echinometra Lucunter Lucunter* (Echinodermata, Echinoidea) in Bermuda. Bull Mar Sci. 82:381–403.

Egger B, Lapraz F, Tomiczek B, Müller S, Dessimoz C, Girstmair J, Škunca N, Rawlinson KA, Cameron CB, Beli E, et al. 2015. A transcriptomic-phylogenomic analysis of the evolutionary relationships of flatworms. Curr Biol. 25:1347–1353.

El-Sayed NM, Myler PJ, Blandin G, Berriman M, Crabtree J, Aggarwal G, Caler E, Renauld H, Worthey EA, Hertz-Fowler C, et al. 2005. Comparative genomics of trypanosomatid parasitic protozoa. Science 309:404–409.

Fairclough SR, Chen Z, Kramer E, Zeng Q, Young S, Robertson HM, Begovic E, Richter DJ, Russ C, Westbrook MJ, et al. 2013. Premetazoan genome evolution and the regulation of cell differentiation in the choanoflagellate Salpingoeca rosetta. Genome Biol. 14: R15.

Figueiredo LM, Rocha EPC, Mancio-Silva L, Prevost C, Hernandez-Verdun D, Scherf A. 2005. The unusually large Plasmodium telomerase reverse-transcriptase localizes in a discrete compartment associated with the nucleolus. Nucleic Acids Res. 33:1111–1122.

Friedman KL, Cech TR. 1999. Essential functions of amino-terminal domains in the yeast telomerase catalytic subunit revealed by selection for viable mutants. Genes Dev. 13:2863–2874.

Giardini MA, Lira CBB, Conte FF, Camillo LR, de Siqueira Neto JL, Ramos CHI, Cano MIN. 2006. The putative telomerase reverse transcriptase component of *Leishmania amazonensis*: gene cloning and characterization. Parasitol Res. 98:447–454.

Gillis AJ, Schuller AP, Skordalakes E. 2008. Structure of the *Tribolium castaneum* telomerase catalytic subunit TERT. Nature 455:633–637.

Gladyshev EA, Arkhipova IR. 2007. Telomere-associated endonuclease-deficient Penelope-like retroelements in diverse eukaryotes. Proc Natl Acad Sci U.S.A. 104:9352–9357.

Gladyshev EA, Arkhipova IR. 2010a. Genome Structure of Bdelloid Rotifers: Shaped by Asexuality or Desiccation? J Hered. 101:S85–S93.

Gladyshev EA, Arkhipova IR. 2010b. A subtelomeric non-LTR retrotransposon Hebe in the bdelloid rotifer Adineta vaga is subject to inactivation by deletions but not 5' truncations. Mob DNA. 1: 12.

Gladyshev EA, Arkhipova IR. 2011. A widespread class of reverse transcriptase-related cellular genes. Proc Natl Acad Sci U.S.A. 108:20311–20316.

González-Estévez C, Felix DA, Smith MD, Paps J, Morley SJ, James V, Sharp TV, Aboobaker AA. 2012. SMG-1 and mTORC1 act antagonistically to regulate response to injury and growth in planarians. PLoS Genet. 8: e1002619.

Greenberg RA, Allsopp RC, Chin L, Morin GB, DePinho RA. 1998. Expression of mouse telomerase reverse transcriptase during development, differentiation and proliferation. Oncogene 16:1723–1730.

Greider CW, Blackburn EH. 1985. Identification of a Specific Telomere Terminal Transferase-Activity in *Tetrahymena* Extracts. Cell 43:405–413.

Greider CW, Blackburn EH. 1987. The telomere terminal transferase of *Tetrahymena* is a ribonucleoprotein enzyme with two kinds of primer specificity. Cell 51:887–898.

Greider CW, Blackburn EH. 1989. A Telomeric sequence in the RNA of *Tetrahymena* Telomerase required for telomere repeat synthesis. Nature 337:331–337.

Hall N, Karras M, Raine JD, Carlton JM, Kooij TWA, Berriman M, Florens L, Janssen CS, Pain A, Christophides GK, et al. 2005. A comprehensive survey of the *Plasmodium* life cycle by genomic, transcriptomic, and proteomic analyses. Science 307:82–86.

Harkisheimer M, Mason M, Shuvaeva E, Skordalakes E. 2013. A motif in the vertebrate telomerase N-terminal linker of TERT contributes to RNA binding and telomerase activity and processivity. Structure 21:1870–1878.

Harley CB, Futcher, AB, Greider CW. 1990. Telomeres shorten during aging of human fibroblasts. Nature 345:458–460.

Harley CB, Villeponteau B. 1995. Telomeres and telomerase in aging and cancer. Curr Opin Genetics Dev. 5:249–255.

Harrington L, Zhou W, McPhail T, Oulton R, Yeung D, Mar V, Bass MB, Robinson MO. 1997. Human telomerase contains evolutionarily conserved catalytic and structural subunits. Genes Dev. 11:3109–3115.

Hashimoto T, Horikawa DD, Saito Y, Kuwahara H, Kozuka-Hata H, Shin-I T, Minakuchi Y, Ohishi K, Motoyama A, Aizu T, et al. 2016. Extremotolerant tardigrade genome and improved radiotolerance of human cultured cells by tardigrade-unique protein. Nat Commun. 7:12808–12814.

Haussmann MF, Winkler DW, Huntington CE, Nisbet ICT, Vleck CM. 2007. Telomerase activity is maintained throughout the lifespan of long-lived birds. Exp Gerontol. 42:610–618.

Hayflick L, Moorhead PS. 1961. The serial cultivation of human diploid cell strains. Exp Cell Res. 25:585–621.

Hedges SB. 2002. The origin and evolution of model organisms. Nat Rev.Genet. 3:838–849.

Hossain S, Singh S, Lue NF. 2002. Functional analysis of the C-terminal extension of telomerase reverse transcriptase. A putative “thumb” domain. J Biol Chem. 277:36174–36180.

Howe KL, Bolt BJ, Cain S, Chan J, Chen WJ, Davis P, Done J, Down T, Gao S, Grove C, et al. 2016. WormBase 2016: expanding to enable helminth genomic research. Nucleic Acids Res. 44:D774–D780.

Hrdličková R, Nehyba J, Bose HR. 2012a. Alternatively Spliced Telomerase Reverse Transcriptase Variants Lacking Telomerase Activity Stimulate Cell Proliferation. Mol Cell Biol. 32:4283–4296.

Hrdličková R, Nehyba J, Lim SL, Grützner F, Bose HR. 2012b. Insights into the evolution of mammalian telomerase: platypus TERT shares similarities with genes of birds and other reptiles and localizes on sex chromosomes. BMC Genomics 13: 216.

Huang H, Chopra R, Verdine GL, Harrison SC. 1998. Structure of a covalently trapped catalytic complex of HIV-1 reverse transcriptase: implications for drug resistance. Science 282:1669–1675.

Huard S, Moriarty TJ, Autexier C. 2003. The C terminus of the human telomerase reverse transcriptase is a determinant of enzyme processivity. Nucleic Acids Res. 31:4059–4070.

Imamura S, Uchiyama J, Koshimizu E, Hanai J-I, Raftopoulou C, Murphey RD, Bayliss PE, Imai Y, Burns CE, Masutomi K, et al. 2008. A Non-Canonical Function of Zebrafish Telomerase Reverse Transcriptase Is Required for Developmental Hematopoiesis. PLoS One 3:e3364–20.

Jaskelioff M, Muller FL, Paik J-H, Thomas E, Jiang S, Adams AC, Sahin E, Kost-Alimova M, Protopopov A, Cadiñanos J, et al. 2011. Telomerase reactivation reverses tissue degeneration in aged telomerase-deficient mice. Nature 469:102–106.

Kao D, Lai AG, Stamataki E, Rosic S, Konstantinides N, Jarvis E, Di Donfrancesco A, Pouchkina-Stancheva N, Semon M, Grillo M, et al. 2016. The genome of the crustacean *Parhyale hawaiensis*, a model for animal development, regeneration, immunity and lignocellulose digestion. eLife 5: e20062.

Katoh K, Asimenos G, Toh H. 2009. Multiple alignment of DNA sequences with MAFFT. Bioinformatics for DNA sequence analysis. 39–64.

Kearse M, Moir R, Wilson A, Stones-Havas S, Cheung M, Sturrock S, Buxton S, Cooper A, Markowitz S, Duran C, et al. 2012. Geneious Basic: an integrated and extendable desktop software platform for the organization and analysis of sequence data. Bioinformatics 28:1647–1649.

Kelleher C, Teixeira MT, Förstemann K, Lingner J. 2002. Telomerase: biochemical considerations for enzyme and substrate. Trends Biochem Sci. 27:1–8.

Kilian A, Bowtell D, Abud HE, Hime GR, Venter DJ, Keese PK, Duncan EL, Reddel RR, Jefferson RA. 1997. Isolation of a candidate human telomerase catalytic subunit gene, which reveals complex splicing patterns in different cell types. Hum Mol Genet. 6:2011–2019.

Kim EB, Fang X, Fushan AA, Huang Z, Lobanov AV, Han L, Marino SM, Sun X, Turanov AA, Yang P, et al. 2011. Genome sequencing reveals insights into physiology and longevity of the naked mole rat. Nature 479:223–227.

Kimura M, Cherkas LF, Kato BS, Demissie S, Hjelmborg JB, Brimacombe M, Cupples A, Hunkin JL, Gardner JP, Lu X, et al. 2008. Offspring's Leukocyte Telomere Length, Paternal Age, and Telomere Elongation in Sperm. PLoS Genet. 4:e37–e39.

Klapper W, Heidorn K, Kuhne K, Parwaresch R, Krupp G. 1998a. Telomerase activity in “immortal” fish. FEBS Lett. 434:409–412.

Klapper W, Kühne K, Singh KK, Heidorn K, Parwaresch R, Krupp G. 1998b. Longevity of lobsters is linked to ubiquitous telomerase expression. FEBS Lett. 439:143–146.

Koutsovoulos G, Kumar S, Laetsch DR, Stevens L, Daub J, Conlon C, Maroon H, Thomas F, Aboobaker AA, Blaxter M. 2016. No evidence for extensive horizontal gene transfer in the genome of the tardigrade *Hypsibius dujardini*. Proc Natl Acad Sci U.S.A. 113:5053–5058.

Kumar S, Hedges SB. 1998. A molecular timescale for vertebrate evolution. Nature 392:917–920.

Lai CK, Mitchell JR, Collins K. 2001. RNA binding domain of telomerase reverse transcriptase. Mol. Cell. Biol. 21:990–1000.

Lai CK, Miller MC, Collins K. 2002. Template boundary definition in *Tetrahymena* telomerase. Genes Dev. 16:415–420.

Lang BF, O’Kelly C, Nerad T, Gray MW, Burger G. 2002. The closest unicellular relatives of animals. Curr Biol. 12:1773–1778.

Lau BW-M, Wong AO-L, Tsao GS-W, So K-F, Yip HK-F. 2008. Molecular cloning and characterization of the zebrafish (*Danio rerio*) telomerase catalytic subunit (telomerase reverse transcriptase, TERT). J Mol Neurosci. 34:63–75.

Li Y, Yates JA, Chen JJL. 2007. Identification and characterization of sea squirt telomerase reverse transcriptase. Gene 400:16–24.

Lingner J, Hughes TR, Shevchenko A, Mann M, Lundblad V, Cech TR. 1997. Reverse transcriptase motifs in the catalytic subunit of telomerase. Science 276:561–567.

Listerman I, Sun J, Gazzaniga FS, Lukas JL, Blackburn EH. 2013. The major reverse transcriptase-incompetent splice variant of the human telomerase protein inhibits telomerase activity but protects from apoptosis. Cancer Res. 73:2817–2828.

Liu SY, Selck C, Friedrich B, Lutz R, Vila-Farré M, Dahl A, Brandl H, Lakshmanaperumal N, Henry I, Rink JC. 2013. Reactivating head regrowth in a regeneration-deficient planarian species. Nature 500:81–84.

Low KC, Tergaonkar V. 2013. Telomerase: central regulator of all of the hallmarks of cancer. Trends Biochem Sci. 38:426–434.

Lue NF. 2004. Adding to the ends: what makes telomerase processive and how important is it? BioEssays 26:955–962.

Malik HS, Burke WD, Eickbush TH. 2000. Putative telomerase catalytic subunits from *Giardia lamblia* and *Caenorhabditis elegans*. Gene 251:101–108.

Martin-Rivera L, Herrera E, Albar JP, Blasco MA. 1998. Expression of mouse telomerase catalytic subunit in embryos and adult tissues. Proc Natl Acad Sci U.S.A. 95:10471–10476.

Mason JM, Randall TA, Capkova Frydrychova R. 2016. Telomerase lost? Chromosoma 125:65–73.

Meier B, Clejan I, Liu Y, Lowden M, Gartner A, Hodgkin J, Ahmed S. 2006. Trt-1 is the *Caenorhabditis elegans* catalytic subunit of telomerase. PLoS Genet. 2:187–197.

Meyerson M, Counter CM, Eaton EN, Ellisen LW. 1997. *hEST2*, the putative human telomerase catalytic subunit gene, is up-regulated in tumor cells and during immortalization. Cell 90:785–795.

Mitchell M, Gillis A, Futahashi M, Fujiwara H, Skordalakes E. 2010. Structural basis for telomerase catalytic subunit TERT binding to RNA template and telomeric DNA. Nat Struct Mol Biol. 17:513–518.

Morand S. 1996. Life-history traits in parasitic nematodes: a comparative approach for the search of invariants. Funct Ecol. 10:210–218.

Moriarty TJ, Huard S, Dupuis S, Autexier C. 2002. Functional multimerization of human telomerase requires an RNA interaction domain in the N terminus of the catalytic subunit. Mol Cell Biol. 22:1253–1265.

Moriarty TJ, Marie-Egyptienne DT, Autexier C. 2004. Functional Organization of Repeat Addition Processivity and DNA Synthesis Determinants in the Human Telomerase Multimer. Mol Cell Biol. 24:3720–3733.

Morrison HG, McArthur AG, Gillin FD, Aley SB, Adam RD, Olsen GJ, Best AA, Cande WZ, Chen F, Cipriano MJ, et al. 2007. Genomic minimalism in the early diverging intestinal parasite *Giardia lamblia*. Science 317:1921–1926.

Muller HJ. 1928. The Production of Mutations by X-Rays. Proc Natl Acad Sci U.S.A. 14:714–726.

Nakamura TM, Morin GB, Chapman KB, Weinrich SL, Andrews WH, Lingner J, Harley CB, Cech TR. 1997. Telomerase catalytic subunit homologs from fission yeast and human. Science 277:955–959.

Nakayama JI, Tahara H, Tahara E, Saito M, Ito K, Nakamura H, Nakanishi T, Ide T, Ishikawa F. 1998. Telomerase activation by hTRT in human normal fibroblasts and hepatocellular carcinomas. Nat Genet. 18:65–68.

Nehyba J, Bose HR, Jr., Hrdličková R. 2012. The Regulation of Telomerase by Alternative Splicing of TERT. INTECH Open Access Publisher.

Nishimura O, Hirao Y, Tarui H, Agata K. 2012. Comparative transcriptome analysis between planarian *Dugesia japonica* and other platyhelminth species. BMC Genomics 13: 289.

Nugent CI, Lundblad V. 1998. The telomerase reverse transcriptase: components and regulation. Genes Dev. 12:1073–1085.

Olovnikov AM. 1973. A theory of marginotomy. The incomplete copying of template margin in enzymic synthesis of polynucleotides and biological significance of the phenomenon. J Theor Biol. 41:181–190.

Owen R, Sarkis S, Bodnar A. 2007. Developmental pattern of telomerase expression in the sand scallop, *Euvola ziczac*. Invertebr Biol. 126:40–45.

O’Reilly M, Teichmann SA, Rhodes D. 1999. Telomerases. Curr Opin Struc Biol. 9:56–65.

Pain A, Renauld H, Berriman M, Murphy L, Yeats CA, Weir W, Kerhornou A, Aslett M, Bishop R, Bouchier C, et al. 2005. Genome of the host-cell transforming parasite *Theileria annulata* compared with *T. parva*. Science 309:131–133.

Pardue ML, DeBaryshe PG. 2003. Retrotransposons Provide an evolutionarily robust non-telomerase mechanism to maintain telomeres. Annu Rev Genet. 37:485–511.

Parkinson EK, Fitchett C, Cereser B. 2008. Dissecting the non-canonical functions of telomerase. Cytogenet Genome Res. 122:273–280.

Peng Y, Mian IS, Lue NF. 2001. Analysis of telomerase processivity: mechanistic similarity to HIV-1 reverse transcriptase and role in telomere maintenance. Mol Cell. 7:1201–1211.

Raible F, Tessmar-Raible K, Osoegawa K, Wincker P, Jubin C, Balavoine G, Ferrier D, Benes V, De Jong P, Weissenbach J, et al. 2005. Vertebrate-type intron-rich genes in the marine annelid *Platynereis dumerilii*. Science 310:1325–1326.

Reznick D, Ghalambor C, Nunney L. 2002. The evolution of senescence in fish. Mech Ageing Dev. 123:773–789.

Riutort M, Field KG, Raff RA, Baguna J. 1993. 18S rRNA sequences and phylogeny of Platyhelminthes. Biochem Sys Ecol. 21:71–77.

Robb SMC, Ross E, Sánchez Alvarado A. 2008. SmedGD: the *Schmidtea mediterranea* genome database. Nucleic Acids Res. 36:D599–D606.

Robb SMC, Gotting K, Ross E, Sánchez Alvarado A. 2015. SmedGD 2.0: The *Schmidtea mediterranea* genome database. Genesis 53:535–546.

Robertson HM. 2009. The choanoflagellate *Monosiga brevicollis* karyotype revealed by the genome sequence: Telomere-linked helicase genes resemble those of some fungi. Chromosome Res. 17:873–882.

Rouda S, Skordalakes E. 2007. Structure of the RNA-Binding Domain of Telomerase: Implications for RNA Recognition and Binding. Structure 15:1403–1412.

Rubio MA, Davalos AR, Campisi J. 2004. Telomere length mediates the effects of telomerase on the cellular response to genotoxic stress. Exp Cell Res. 298:17–27.

Rudolph KL, Chang S, Lee H-W, Blasco M, Gottlieb GJ, Greider C, DePinho RA. 1999. Longevity, stress response, and cancer in aging telomerase-deficient mice. Cell 96:701–712.

Ruiz-Trillo I, Roger AJ, Burger G, Gray MW, Lang BF. 2008. A Phylogenomic Investigation into the Origin of Metazoa. Mol Biol Evol. 25:664–672.

Ryan JF, Pang K, Schnitzler CE, Nguyen AD, Moreland RT, Simmons DK, Koch BJ, Francis WR, Havlak P, NISC Comparative Sequencing Program, et al. 2013. The Genome of the ctenophore *Mnemiopsis leidyi* and its implications for cell type evolution. Science 342:1242592–1242592.

Samper E, Flores JA, Blasco MA. 2001. Restoration of telomerase activity rescues chromosomal instability and premature aging in Terc(-/-) mice with short telomeres. EMBO rep. 2:800–807.

Schumpert C, Nelson J, Kim E, Dudycha JL, Patel RC. 2015. Telomerase Activity and Telomere Length in *Daphnia*. PLoS ONE 10: e0127196.

Sekaran VG, Soares J, Jarstfer MB. 2010. Structures of telomerase subunits provide functional insights. BBA - Proteins and Proteom. 1804:1190–1201.

Shalchian-Tabrizi K, Minge MA, Espelund M, Orr R, Ruden T, Jakobsen KS, Cavalier-Smith T. 2008. Multigene phylogeny of choanozoa and the origin of animals. PLoS ONE 3:e2098–7.

Smith LL, Coller HA, Roberts JM. 2003. Telomerase modulates expression of growth-controlling genes and enhances cell proliferation. Nat Cell Biol. 5:474–479.

Srivastava M, Simakov O, Chapman J, Fahey B, Gauthier MEA, Mitros T, Richards GS, Conaco C, Dacre M, Hellsten U, et al. 2010. The *Amphimedon queenslandica* genome and the evolution of animal complexity. Nature 466:720–726.

Stamatakis A. 2014. RAxML version 8: a tool for phylogenetic analysis and post-analysis of large phylogenies. Bioinformatics. 30:1312–1313.

Steele RE, David CN, Technau U. 2011. A genomic view of 500 million years of cnidarian evolution. Trends Genet. 27:7–13.

Steenstrup T, Hjelmborg JVB, Kark JD, Christensen K, Aviv A. 2013. The telomere lengthening conundrum artifact or biology? Nucleic Acids Res. 41:e131–e131.

Stevens JR, Noyes HA, Schofield CJ, Gibson W. 2001. The molecular evolution of Trypanosomatidae. Adv parasitol. 48:1–53.

Stone RC, Horvath K, Kark JD, Susser E, Tishkoff SA, Aviv A. 2016. Telomere length and the cancer-atherosclerosis trade-off. PLoS Genet. 12:e1006144–10.

Suram A, Herbig U. 2014. The replicometer is broken: telomeres activate cellular senescence in response to genotoxic stresses. Aging Cell 13:780–786.

Sýkorová E, Fajkus J. 2009. Structure-function relationships in telomerase genes. Biol Cell. 101:375–392.

Szostak JW, Blackburn EH. 1982. Cloning Yeast Telomeres on Linear Plasmid Vectors. Cell 29:245–255.

Sæbøe-Larssen S, Fossberg E, Gaudernack G. 2006. Characterization of novel alternative splicing sites in human telomerase reverse transcriptase (hTERT): analysis of expression and mutual correlation in mRNA isoforms from normal and tumour tissues. BMC Mol Biol. 7: 26.

Tan TCJ, Rahman R, Jaber-Hijazi F, Felix DA, Chen C, Louis EJ, Aboobaker A. 2012. Telomere maintenance and telomerase activity are differentially regulated in asexual and sexual worms. Proc Natl Acad Sci U.S.A. 109:4209–4214.

Tasaka K, Yokoyama N, Nodono H, Hoshi M, Matsumoto M. 2013. Innate sexuality determines the mechanisms of telomere maintenance. Int J Dev Biol. 57:69–72.

Tomás-Loba A, Flores I, Fernández-Marcos PJ, Cayuela ML, Maraver A, Tejera A, Borrás C, Matheu A, Klatt P, Flores JM, et al. 2008. Telomerase Reverse Transcriptase Delays Aging in Cancer-Resistant Mice. Cell 135:609–622.

Tomlinson CG, Holien JK, Mathias JAT, Parker MW, Bryan TM. 2016. The C-terminal extension of human telomerase reverse transcriptase is necessary for high affinity binding to telomeric DNA. Biochimie 128-129:114–121.

Torruella G, Derelle R, Paps J, Lang BF, Roger AJ, Shalchian-Tabrizi K, Ruiz-Trillo I. 2012. Phylogenetic Relationships within the Opisthokonta Based on Phylogenomic Analyses of Conserved Single-Copy Protein Domains. Mol Biol Evol. 29:531–544.

Tsai IJ, Zarowiecki M, Holroyd N, Garciarrubio A, Sanchez-Flores A, Brooks KL, Tracey A, Bobes RJ, Fragoso G, Sciutto E, et al. 2013. The genomes of four tapeworm species reveal adaptations to parasitism. Nature 496:57–63.

Tzfati Y, Fulton TB, Roy J, Blackburn EH. 2000. Template boundary in a yeast telomerase specified by RNA structure. Science 288:863–867.

Ulaner GA, Hu JF, Vu TH, Giudice LC, Hoffman AR. 1998. Telomerase activity in human development is regulated by human telomerase reverse transcriptase (hTERT) transcription and by alternate splicing of hTERT transcripts. Cancer Res. 58:4168–4172.

Ulaner GA, Hu JF, Vu TH, Oruganti H, Giudice LC, Hoffman AR. 2000. Regulation of telomerase by alternate splicing of human telomerase reverse transcriptase (hTERT) in normal and neoplastic ovary, endometrium and myometrium. Int J Cancer 85:330–335.

Vaziri H, Dragowska W, Allsopp RC, Thomas TE, Harley CB, Lansdorp PM. 1994. Evidence for a Mitotic Clock in Human Hematopoietic Stem-Cells - Loss of Telomeric Dna with Age. Proc Natl Acad Sci U.S.A. 91:9857–9860.

Wasik K, Gurtowski J, Zhou X, Ramos OM, Delás MJ, Battistoni G, Demerdash El O, Falciatori I, Vizoso DB, Smith AD, et al. 2015. Genome and transcriptome of the regeneration-competent flatworm, *Macrostomum lignano*. Proc Natl Acad Sci U.S.A. 112:12462–12467.

Watson JD. 1972. Origin of concatemeric T7 DNA. Nature New Biol. 239:197–201.

Welch DBM, Meselson M. 2000. Evidence for the evolution of bdelloid rotifers without sexual reproduction or genetic exchange. Science 288:1211–1215.

Wheeler NJ, Agbedanu PN, Kimber MJ, Ribeiro P, Day TA, Zamanian M. 2015. Functional analysis of Girardia tigrina transcriptome seeds pipeline for anthelmintic target discovery. Parasit Vectors. 8: 34.

Withers JB, Ashvetiya T, Beemon KL. 2012. Exclusion of exon 2 Is a common mRNA splice variant of primate telomerase reverse transcriptases. PLoS ONE 7:e48016–e48019.

Wong MS, Chen L, Foster C, Kainthla R, Shay JW, Wright WE. 2013. Regulation of Telomerase Alternative Splicing: A Target for Chemotherapy. Cell Reports 3:1028–1035.

Wong MS, Wright WE, Shay JW. 2014. Alternative splicing regulation of telomerase: a new paradigm? Trends Genet. 30:430–438.

Wurm Y, Wang J, Riba-Grognuz O, Corona M, Nygaard S, Hunt BG, Ingram KK, Falquet L, Nipitwattanaphon M, Gotzek D, et al. 2011. The genome of the fire ant *Solenopsis invicta*. Proc Natl Acad Sci U.S.A. 108:5679–5684.

Wyatt HDM, West SC, Beattie TL. 2010. InTERTpreting telomerase structure and function. Nucleic Acids Res. 38:5609–5622.

Xu P, Widmer G, Wang YP, Ozaki LS, Alves JM, Serrano MG, Puiu D, Manque P, Akiyoshi D, Mackey AJ, et al. 2004. The genome of *Cryptosporidium hominis*. Nature 431:1107–1112.

Yang Y, Xiong J, Zhou Z, Huo F, Miao W, Ran C, Liu Y, Zhang J, Feng J, Wang M, et al. 2014. The genome of the myxosporean *Thelohanellus kitauei* shows adaptations to nutrient acquisition within its fish host. Genome Biol Evol. 6:3182–3198.

Young ND, Nagarajan N, Lin SJ, Korhonen PK, Jex AR, Hall RS, Safavi-Hemami H, Kaewkong W, Bertrand D, Gao S, et al. 2014. The *Opisthorchis viverrini* genome provides insights into life in the bile duct. Nat Commun. 5: 4378.

